# Improving splice site usage prediction with SPLAIRE

**DOI:** 10.64898/2026.06.08.731019

**Authors:** Matthew Runyan, Saumya Gupta, Yul Leshaem, David Geller-McGrath, Congjian Liu, Aabida Saferali, Jennifer Dy, Predrag Radivojac, Yohannes Tesfaigzi, Peter Castaldi, Ayan Paul

## Abstract

**Background:** Alternative splicing, the mechanism by which intronic sequences are excised from pre-mRNAs to produce mature mRNA, affects >95% of human protein-coding genes and is a major driver of human disease states. The spliceosome, a protein-RNA complex responsible for splicing pre-mRNA, identifies candidate splice sites partly through the recognition of characteristic sequence motifs at exon-intron junctions. Deep learning models that predict the presence of splice sites from pre-mRNA sequence have achieved breakthrough performance relative to previous machine-learning techniques, and these models have improved our ability to identify pathogenic genetic variants that alter splicing.

**Results:** We show that, while overall performance measures from these models suggest near-perfect performance, substantial gaps in prediction remain, including the identification of splice sites with low usage rates and tissue-specific splice sites. We leverage one of the largest paired RNA and genotyping datasets used to date to train a novel splicing model optimized for a specific cell type, human airway epithelial cells. We trained a dilated convolutional neural network on data from cultured airway epithelial cells from 100 donors, and showed that this model outperforms current state-of-the-art models on splice site identification and splice site usage quantification, including on multiple tissues not included in the model training data.

**Conclusions:** We present the most comprehensive evaluation of state-of-the-art splicing models published to date, revealing reasonable performance across models for genetic variant effect prediction along with important performance gaps and insights into directions for future model development.

## 1 Background

Nearly all human genes undergo alternative splicing which is a major driver of isoform and proteomic diversity. Amongst the several important components of cellular processes that drive diversity within and between organisms, alternative splicing plays an important role in establishing cell identity, tissue function, and disease phenotypes [1–5]. While splice site usage is readily quantified through RNA sequencing (RNA-seq), there is substantial interest in the development of sequence-based splicing models that can be used to identify genetic variants that will affect splicing. One important use case of these models is in the area of variant effect prediction for genetic testing [6, 7], because a substantial proportion of human disease is caused by genetically-altered splicing [8, 9].

One of the first computational models of splicing, MaxEntScan, analyzed raw sequence in the immediate vicinity of the splice site, demonstrating the capacity of computational approaches to recognize core motifs responsible for creating splice sites [10]. Subsequent models achieved high overall accuracy for various splicing outcomes by relying on “hand-engineered” features that captured important aspects of splicing such exon structure and RBP-binding [11, 12]. The application of deep learning to splicing, combined with a return to the compositional analysis of longer stretches of sequence, led to a breakthrough in predictive performance [13–16] using dilated convolutional neural networks [17, 18] (CNN) with residual connections [19] and Transformers. The very high values of performance metrics reported in these papers demonstrated the performance gains made by these models, but independent evaluations more representative of real world use also demonstrated the limitations of these models [20–22].

Recently, foundation models have been built for DNA and RNA sequences which can be used for predicting alternative splicing; however when models such as HyenaDNA [23], SegmentNT [24], DNABERT [25, 26], and Evo [27, 28] are evaluated on splice site prediction they tend to underperform state-of-the-art dedicated splicing models such as SpliceAI [14], Pangolin [20], or recent sequence-to-profile models [29, 30]. Moreover, foundation models have high parameteric complexity with hundreds of millions or billions of parameters, making them difficult to train and unwieldy for any downstream task that requires fine tuning. The dilated CNN based models that we discuss in this paper have a parametric complexity of under or about a few million parameters and are fast for training and inference, leading to more flexible application in real world settings and more convenient implementation of interpretability measures to query the model for important sequence features. Other recent approaches include OpenSpliceAI [31], which provides a modular reimplementation enabling retraining across species, and reference-informed models trained across multiple mammals [32].

The goal of this project was to develop a state-of-the-art sequence-based splicing model by training on one of the largest paired genotyping and RNA-seq datasets used to date, leveraging data from cultured human airway epithelial cells (HAECs) from >100 independent donors. We hypothesized that training a dilated CNN on 100 samples using actual genotype information would lead to superior performance in splice site identification, quantification, and variant effect prediction. By using dual task prediction of binary splice site presence and continuous splice site usage (SSU), we hypothesized that our model would be better able to learn distinct sequence patterns associated with high and low-usage splice sites. To evaluate our model, we compared it’s performance in multiple domains against three state-of-the-art sequence based splicing models - SpliceAI, Pangolin, and SpliceTransformer [21]. With this approach, our models achieved state-of-the-art performance on HAEC data and other tissues from GTEx, substantially outperforming other models on SSU prediction across multiple tissues. For variant effect prediction, we observed that all evaluated models performed within a relatively tight range with reasonable ability to identify causal sQTL variants under distance-matched evaluation (AUPRC 0.76–0.80 across 49 GTEx tissues at PIP *>* 0.9). In a detailed evaluation on tissue-specific splicing performance, we observed that the two models with tissue-specific splicing heads, Pangolin and SpliceTransformer, have substantial gaps in their ability to predict tissue-specific splice events. These results show the utility of existing state-of-the-art splicing models while highlighting specific areas of variant effect detection and tissue-specific performance that require further improvement.

### 1.1 Training Data

We compare our model to three models that are the current state-of-the-art. In this section, we briefly review the datasets used to train these models.

#### SpliceAI

was trained on a subset of human chromosomes from GENCODE-annotated pre-mRNA transcript sequences [33]. Transcripts on the remaining chromosomes, with paralogs excluded, were used to test predictions. SpliceAI has only a single prediction head that is trained on binary labels (splice site or not a splice site), so it should be noted that SSU predicted by SpliceAI or any scores derived from it assume that the probability of a nucleotide being a splice site is interpreted as the usage of that splice site.

#### Pangolin

was trained using data from four species – human, rhesus macaque, rat, and mouse – sourced from [34]. Sequence and RNA splicing measurements were processed using SpliSER [35] to quantify the usage of all splice sites after mapping RNA-seq data from heart, liver, brain, and testis from up to 8 samples per species per tissue. The training set consisted of genes on human chromosomes 2, 4, 6, 8, 10–22, X, and Y, including splice and non-splice sites, while the test set consisted of positions from genes on human chromosomes 1, 3, 5, 7, and 9. When training on the data from rhesus macaque, rat, and mouse, intra-species orthologs or paralogs of genes from the test human chromosomes were excluded. Unlike SpliceAI, Pangolin architecture included two types prediction heads, one for splice site classification (trained on binary splice site labels) and another for continuous SSU prediction (trained on SSUs calculated from RNA-seq data using Spliser). Pangolin outputs four tissue-specific predictions.

#### SpliceTransformer

was trained on GTEx V8 [36] and the same multi-organism RNA-seq dataset [34] used by Pangolin. Like Pangolin, SpliceTransformer had a tissue classification and 15 tissue-specific SSU prediction heads, but the authors took the approach of quantifying SSU not at the level of individual samples (direct SSU quantification such as with Spliser) but rather by representing SSU as the proportion of samples belonging to the tissue that contained corresponding splice junctions. For the data from Ref. [34], splicing quantifications were derived using SpliSER [35] as was done for the data used to train Pangolin. Gene sequences that have paralogs were excluded from the testing dataset.

We use the following datasets for training our models and for comparing with the performance of the other models:

#### The SPLicing in AIRway epithelium (SPLAIRE)

model was trained on RNA-seq data generated from 100 primary HAECs cultured in basal media. Like Pangolin and SpliceTransformer, SPLAIRE had a classification and a quantification head that were trained on binary splice site labels and Spliser-generated SSU values, respectively. A separate model, SPLAIRE-Var was also trained on matching genotype data such that the input sequences included correct genotype data for each individual (rather than the reference genome). We also selected 10 samples for use as testing data and identified splice sites as SSU values using the same methods.

### Tissue-specific gene expression and splicing in GTEx

The GTEx project [36] provides paired RNA-seq and genotyping data across multiple human tissues, supporting cross-tissue evaluation of splicing models. We selected 10 donors from the GTEx v10 release with paired RNA-seq and Whole Genome Sequencing (WGS) data in four tissues (lung, whole blood, brain cortex, and testis). RNA-seq alignments and SSU values were processed using the same pipeline applied to HAEC data (Methods). These GTEx data are used as a held-out evaluation set and were not included in any training sets used for the SPLAIRE models.

## 2 Results

### 2.1 Splice site usage across tissues

We quantified splice site usage (SSU) from short-read RNA-seq in human airway epithelial cells (HAEC) and four GTEx tissues (lung, testis, brain cortex, and whole blood). SSU was computed from 10 held-out donors per tissue (same GTEx donors across the four tissues; Methods). SSU represents the proportion of reads supporting a splice event at a given position, ranging from 0 (never used) to 1 (constitutively used). The overlap of splice sites detected across tissues is shown in Figure 2a. Testis contained the most splice sites, consistent with the high levels of transcriptional activity known to occur in this tissue. The distribution of mean SSU across samples was bimodal in all tissues, with peaks at low and high usage (Figure 2b). Among splice sites detected in each tissue that also appear in MANE Select transcripts, SSU distributions were strongly skewed toward 1, indicating near-constitutive usage across all five tissues (Figure 2c).

**Figure 1.**
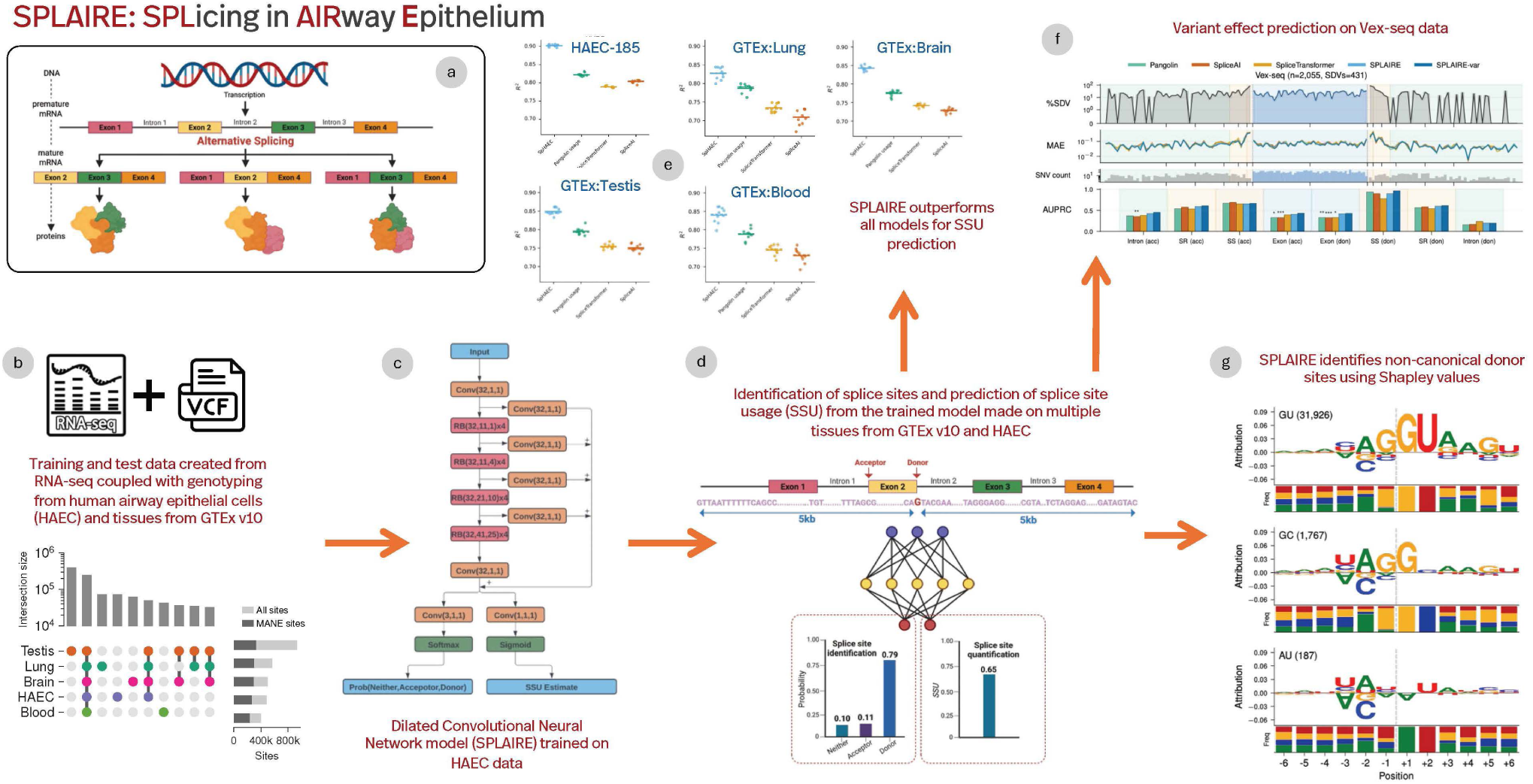
An overview of the development of and analysis using SPLAIRE. Panel (**a**) provides an overview of the splicing mechanism, emphasizing on how it contributes to proteome diversity. Panel (**b**) defines the training and inference data used for SPLAIRE. RNA sequencing along with genome-wide genotyping was used to create the datasets from human airway epithelial cell (HAEC) and several tissues from GTEx. In Panel (**c**), we show the architecture of the ML model being used. The dilated convolutional neural network with residual connections is derived from the SpliceAI architecture, with the difference that there are two prediction heads, one for classification and the other for regression. Panel (**d**) defines the input and the output of the ML models, with one-hot encoded RNA sequence being provided as an input, the output being the classification of the central nucleotide as an acceptor, donor, or neither, and a prediction of the splice site usage (SSU). SSU is computed using SpliSER. Panel (**e**) and (**f**) highlight some of the results derived from the predictions. In the former, we compare against other state-of-the-art ML models for splicing. In the latter, we study variant effects using reporter assays. In panel (**g**) we highlight some of the interpretations that we perform using model predictions and Shapley values. We see that the model we train uses known splicing motifs to determine a splice site and its usage.

**Figure 2.**
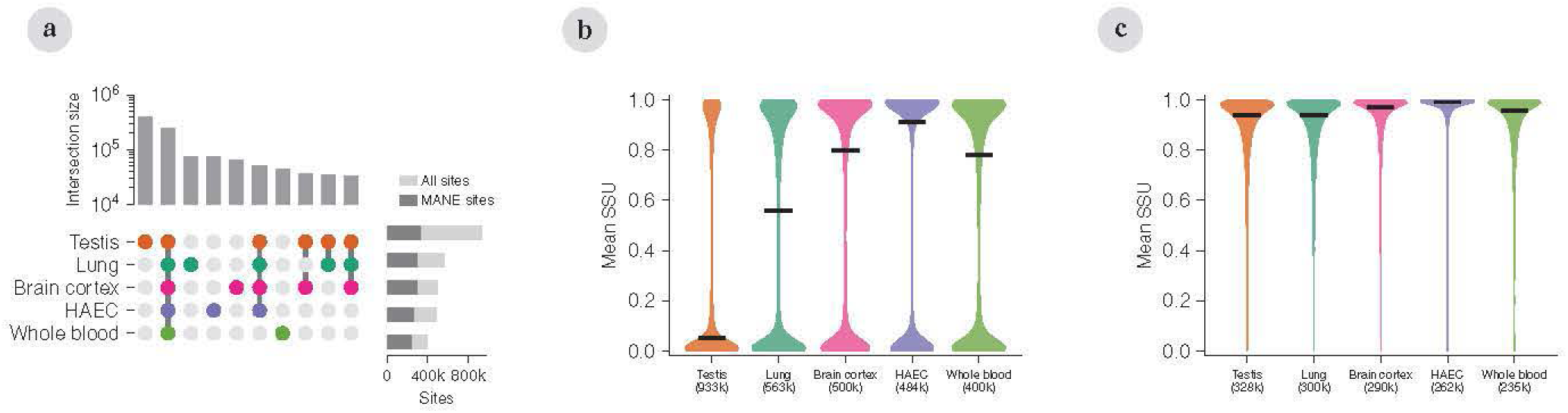
Overview of splice site datasets. Splice sites were quantified from human airway epithelial cells (HAEC, 10 held-out samples) and four GTEx tissues (lung, testis, brain cortex, and whole blood, 10 samples each from the same 10 individuals) across all autosomal protein-coding genes. Splice site usage (SSU) is the fraction of reads at a given splice site that support splicing at that site, ranging from 0 (never used) to 1 (constitutively used). (**a**) UpSet plot showing the overlap of splice sites across the five tissues. The top bar chart shows intersection sizes on a log scale. The right horizontal bars show total sites detected per tissue, with the subset present in MANE Select transcripts in dark grey. (**b**) Violin plot of mean SSU (across held-out individuals) for all detected splice sites in each tissue, with the horizontal line within each violin marking the median. The bimodal distribution with peaks at low and high usage is visible in each tissue. (**c**) Violin plot of mean SSU restricted to splice sites present in MANE Select transcripts. The distribution is strongly skewed toward high SSU compared with panel **b**.

### 2.2 Model Classification and Regression Evaluation

We compared SPLAIRE against SpliceAI, Pangolin, and SpliceTransformer on held-out test genes from non-paralogous protein-coding genes on chromosomes 1, 3, 5, and 7 across all five tissues (HAEC, lung, testis, brain cortex, and whole blood, 10 held-out donors each). Our evaluations covered classification performance (differentiating splice sites from non-splice sites), regression performance (correlation of predicted SSU to actual SSU at splice sites), and tissue-specific prediction.

For classification performance, we treated annotated splice sites as positives and all other nucleotide positions as negatives, and we evaluated against three sets of splice sites: 1) all GENCODE v45 splice sites contained in protein coding transcripts, 2) a more restrictive set using only splice sites from MANE Select transcripts [37], and 3) all splice sites observed in the RNA-seq data, including unannotated and low-usage sites absent from reference annotations. Between 28% and 51% of RNA-seq-observed sites in each tissue were absent from GENCODE, with testis having the largest number of unannotated sites. On set 1, SpliceAI achieved the highest AUPRC (0.96), followed by SPLAIRE (0.95; Table 6). On set 2, SPLAIRE achieved the highest AUPRC (0.95) while SpliceAI dropped to 0.87. The drop in performance for SpliceAI between set 1 and 2 may be due to the fact that this model was trained directly on the GENCODE annotations comprising set 1. On set 3, all splice sites detected in our RNA-seq data, AUPRC across all models dropped to 0.68–0.77. SPLAIRE achieved the highest AUPRC in four of five tissues (0.74–0.77), with SpliceAI and SPLAIRE both achieving AUPRC of 0.68 on testis (Figure 3a,c).

**Figure 3.**
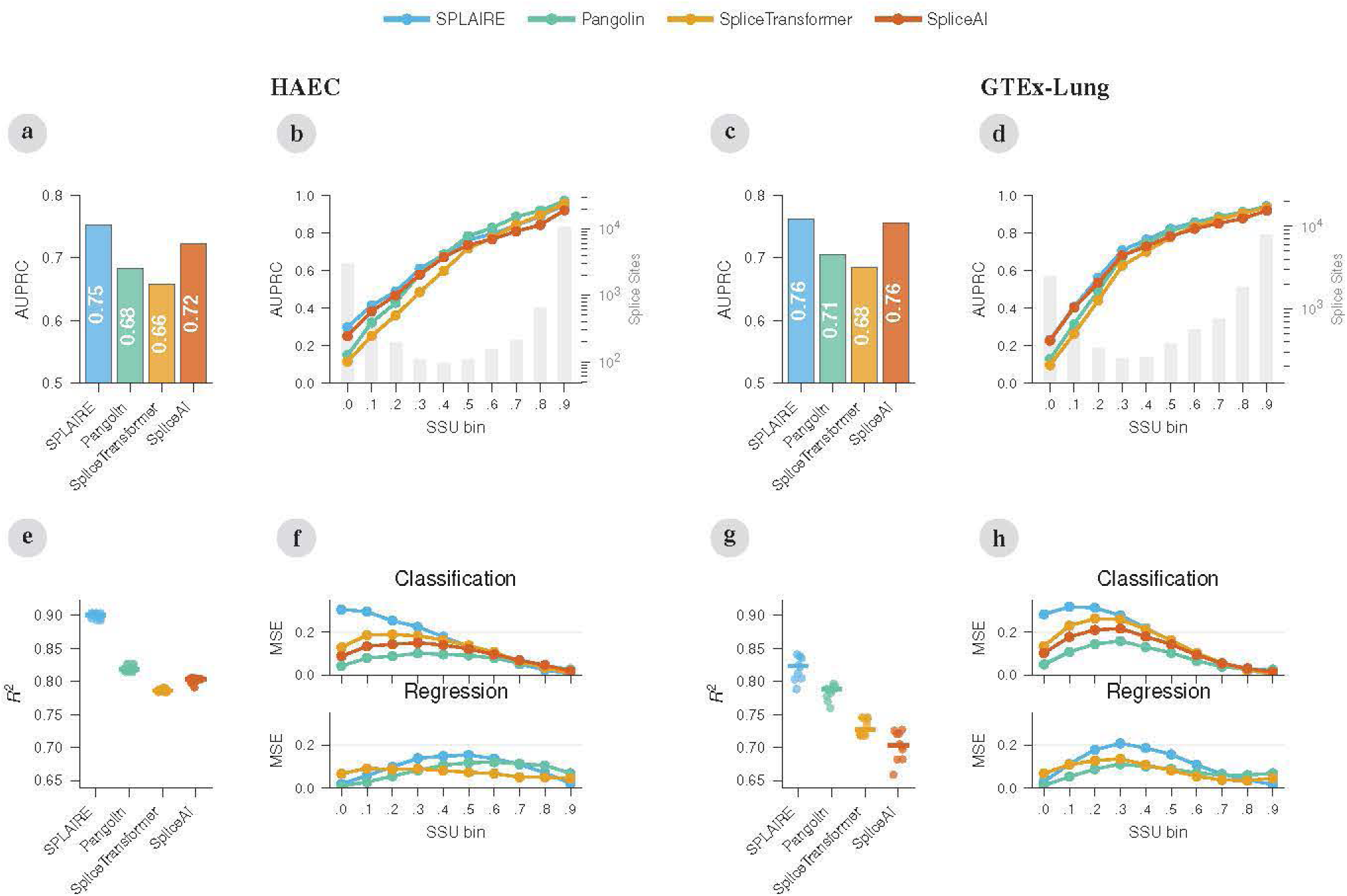
Splice site classification and SSU prediction across tissues. Performance of SPLAIRE, SpliceAI, Pangolin, and SpliceTransformer. Pangolin and SpliceTransformer predictions are averaged across tissue heads. For each tissue (HAEC and GTEx-Lung, 10 held-out donors each), the same four panels are shown. (**a**,**c**) Splice site classification AUPRC for splice sites observed in RNA-seq. (**b**,**d**) AUPRC stratified by SSU decile. The secondary axis shows the number of splice sites per bin. (**e**,**g**) *R*^2^ for SSU prediction, where each point is one held-out individual and the line shows the median. (**f**,**h**) Mean squared error for SSU prediction stratified by SSU decile. Top row shows classification, bottom row shows regression.

For regression performance, we evaluated the heads from the models that had been trained on quantiative splice site usage (SSU) rather than binary labels. Since SpliceAI was only trained on binary labels, we used the maximum of its acceptor and donor classification probabilities as an SSU proxy. SPLAIRE achieved the highest *R*^2^ across all five tissues (0.82–0.90), followed by Pangolin (0.78–0.82), SpliceTransformer (0.73–0.79), and SpliceAI (0.70–0.80; Figure 3e,g). SPLAIRE achieved *R*^2^ = 0.90 on HAEC, the tissue on which it was trained, and maintained strong performance on GTEx tissues (lung *R*^2^ = 0.82, brain cortex *R*^2^ = 0.84, whole blood *R*^2^ = 0.83). We also evaluated all models on Pangolin’s test set across four tissues (heart, liver, brain, testis), where SPLAIRE achieved the highest *R*^2^ (0.62–0.68), outperforming Pangolin (*R*^2^ = 0.31–0.51; Supplementary Figure S3, Supplementary Table 9). We examined model calibration by regressing measured SSU on predicted SSU per model and tissue (Supplementary Figure S2). SPLAIRE’s linear fit showed the best calibration. Pangolin underpredicted across the full SSU range, and notably its classification head achieved higher *R*^2^ than its regression head on every tissue (0.80–0.87 vs 0.77–0.82, Supplementary Table 7). SpliceAI consistently overpredicted the true SSU.

We observed that prediction performance decreased substantially for “weaker” splice sites with lower SSU. Stratifying splice sites into deciles of mean SSU, classification AUPRC declined substantially with decreasing usage for all models (Figure 3b,d). At the lowest decile (SSU 0–0.1), SPLAIRE and SpliceAI were the top performers in both tissues (HAEC, SPLAIRE 0.30 and SpliceAI 0.25; lung, SPLAIRE 0.23 and SpliceAI 0.23), while Pangolin and SpliceTransformer fell below 0.15. Regression MSE showed the opposite pattern, with lowest error at the SSU extremes (near 0 and 1) and highest error at intermediate deciles (Figure 3f,h). Top-line classification performance is therefore strong in datasets dominated by high-usage sites, but all models struggle to differentiate low-usage splice sites from non-splice positions and to predict intermediate SSU accurately.

To evaluate regression behavior on annotated versus unannotated splice junctions in set 3, we computed metrics separately on the two subsets (Methods). On annotated sites, *R*^2^ was 0.63–0.77 for SPLAIRE and lower for the other models (SpliceAI 0.53–0.69, Pangolin 0.44–0.54, SpliceTransformer 0.52–0.60). On unannotated sites, *R*^2^ was negative for all models because true SSU values concentrate near zero with very small variance, making *R*^2^ uninformative on this subset (Supplementary Figure S4). We therefore report MAE for this subset, which remained small for all models (SPLAIRE 0.0015–0.0022, Pangolin 0.0020–0.0031, SpliceAI 0.030–0.062, SpliceTransformer 0.032–0.049; Supplementary Table 4) because both true SSU and predictions cluster near zero.

Pangolin and SpliceTransformer were trained to make tissue-specific splicing predictions. To quantify how well these models performed on tissue-specific splice sites, we computed the tissue specificity index *τ* [38], which ranges from 0 (uniform usage across tissues) to 1 (usage concentrated in one tissue), and its reverse *τ*_rev_, which ranges from 0 to 1 in the opposite direction (high when one tissue has low usage while others remain high) (Methods). We applied these indices to splice sites from Pangolin’s test set (four tissues with 36,727 sites detected) where tissue-matched heads are available for Pangolin and SpliceTransformer. We grouped sites into three categories - constitutive sites (*τ <* 0.1, *n* = 26,121) have near-zero specificity, intermediate sites (0.1 ≤ *τ <* ≤ 0.5, *n* = 6,995), and tissue-specific sites (*τ* ≥ 0.5 or *τ*_rev_ ≥ 0.7, additionally requiring cross-tissue SSU range ≥ 0.3 to exclude low-signal sites, *n* = 1,746, Supplementary Figure S5). Tissue-specific sites were further classified into inclusion (high SSU in the target tissue) and exclusion (low SSU). Testis had the most tissue-specific splice sites (Figure 4a), and ΔSSU (the difference between the target tissue and the mean of the remaining tissues) was distributed as expected (Figure 4b). Raw SSU of the target tissue confirmed that inclusion sites have high absolute usage while exclusion sites have near-zero usage in the target tissue (Figure 4c). We observed that for tissue-specific sites, the performance of the tissue-matched heads of both Pangolin and SpliceTransformer was usually worse than the performance of at least one of the other heads from the same model for a different tissue (Figure 4d), demonstrating lack of tissue-specificity for model predictions on this set of splice sites with clear tissue-specific behavior. To investigate whether tissue-matched models learn distinct sequence features at tissue-specific sites, we compared DeepLIFT-SHAP attributions across all four Pangolin models at brain-specific splice sites. Attribution patterns were nearly identical across models (median pairwise Pearson *r* = 0.99 for both normalized and raw attributions, Supplementary Figure S6), indicating that all four models identify the same sequence features and assign them the same relative importance despite producing different predictions. Tissue specificity in Pangolin therefore appears to be encoded almost entirely in the final 1 × 1 convolution mapping a shared feature representation to per-tissue SSU, rather than in the upstream sequence features themselves.

**Figure 4.**
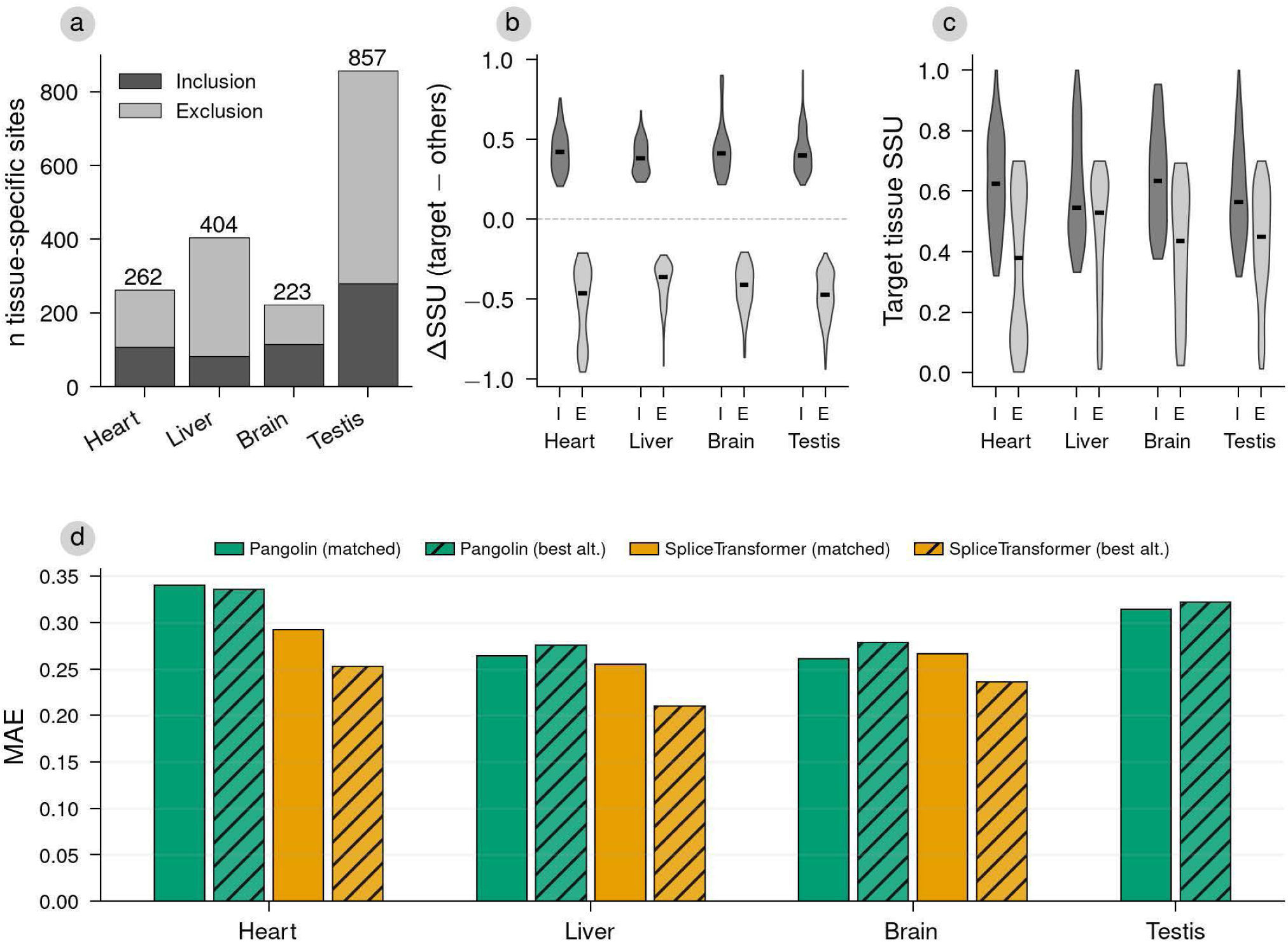
Tissue specificity characterization and model performance. Splice sites detected in the four tissues in Pangolin’s test set (heart, liver, brain, testis, 36,727 sites) were classified using *τ* and *τ*_rev_ into three groups. Constitutive sites (*n* = 26,121), intermediate sites (*n* = 6,995), and tissue-specific sites (*τ* ≥ 0.5 or *τ*_rev_ ≥ 0.7, cross-tissue SSU range ≥ 0.3, *n* = 1,746). (**a**) Inclusion and exclusion direction counts per tissue. At each tissue-specific site, the target tissue (the tissue whose mean SSU deviated most from the cross-tissue mean) was classified as inclusion-specific (highest SSU) or exclusion-specific (lowest SSU). (**b**) ΔSSU (target tissue SSU minus mean of remaining tissues) split by direction per tissue. I = inclusion, E = exclusion. (**c**) SSU of the target tissue, split by direction per tissue. (**d**) MAE on tissue-specific sites per target tissue. Solid bars show the tissue-matched head and hatched bars show the best alternative head (lowest MAE among non-matched heads). SPLAIRE and SpliceAI do not have tissue-specific heads and are not shown, while SpliceTransformer is omitted for testis as it does not have a testis-specific head.

### 2.3 Reporter Assay Evaluation

One of the main use cases for sequence-based splicing models is the identification of functional genetic variants, an important problem in clinical genetic testing. We evaluated the prediction of variant effects in two massively parallel reporter assays (MPRAs) [40] that measure variant effects on exon inclusion in *in vitro* minigene contexts. Vex-seq [41] tests 2,055 variants (1,960 SNVs and 95 indels) across 110 alternatively spliced exons using a barcode-based sequencing readout in HepG2 cells. MFASS [39] evaluates 27,733 SNVs identified in the Exome Aggregation Consortium across 2,198 constitutive exons using a fluorescence-based Sort-seq readout in HEK293T cells. We also evaluated SPLAIRE-var, an identical model to SPLAIRE but trained on pre-mRNA sequences containing personal genetic variants rather than the reference genome. For each variant we computed the alt-minus-ref delta at the acceptor and donor splice sites of the tested exon and averaged the two deltas to obtain the predicted ΔPSI (Percent Spliced-In). On Vex-seq, SPLAIRE-var achieved the highest Pearson correlation and AUPRC (0.60, 0.36), followed by SPLAIRE (0.59, 0.35; Figure 5a,b). SpliceAI (0.55, 0.31), Pangolin (0.55, 0.30), and SpliceTransformer (0.53, 0.33). On MFASS, Pangolin achieved the highest Pearson correlation and AUPRC (0.54, 0.35), followed by SpliceAI (0.52, 0.34), SPLAIRE (0.49, 0.33), and SpliceTransformer (0.46, 0.30; Figure 5c,d). Despite being trained on sequences containing personal genetic variants, SPLAIRE-var showed only marginal improvement over SPLAIRE on Vex-seq (Pearson 0.60 vs 0.59) and no improvement on MFASS.

**Figure 5.**
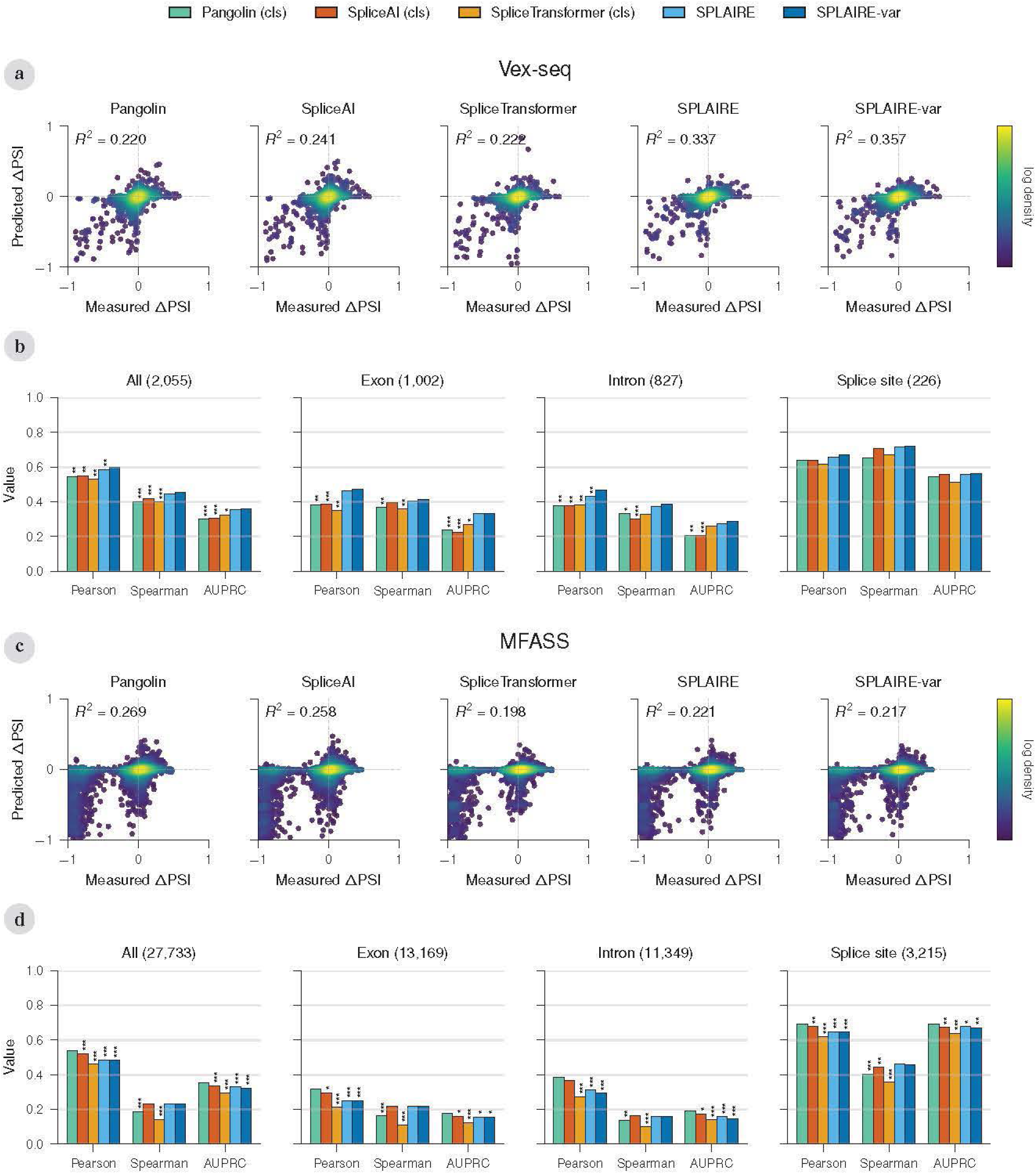
Variant effect prediction on reporter assays. **(a)** Scatter plots of measured versus predicted ΔPSI for each model on Vex-seq (n = 2,055 variants, 110 exons), colored by log density. **(b)** Pearson correlation, Spearman correlation, and AUPRC on Vex-seq stratified by variant location. Splice site proximal (3 exonic and 8 intronic nucleotides adjacent to each splice junction [39], n = 226), Exon (n = 1,002), Intron (n = 827). Exon and Intron categories exclude splice site proximal variants. Pearson and Spearman computed on all variants within each location. AUPRC computed treating splice-disrupting variants (|ΔPSI| > 0.10 for Vex-seq, > 0.50 for MFASS) as positives. Significance stars denote pairwise comparisons against the best-performing model (paired bootstrap, 10,000 resamples, Holm–Bonferroni corrected). **(c,d)** Same as (a–b) for MFASS (n = 27,733 variants, 2,198 exons; splice site n = 3,215, exon n = 13,169, intron n = 11,349).

Stratifying variants by position relative to the tested exon (Figure 5b,d) showed that correlations and AUPRC were highest for splice site proximal variants (i.e. variants located within 3 exonic and 8 intronic nucleotides of each splice junction; *n* = 226 on Vex-seq, *n* = 3,215 on MFASS) and dropped substantially for exonic and intronic variants. As expected, the ΔPSI values were also substantially larger for splice site proximal variants than variants located in distal exonic and intronic positions. Per-position error profiles were not obviously distinguishable between models in both assays, demonstrating that the performance of these models and their error modes were similar. All models overpredicted effects for neutral variants (measured ΔPSI within 1 SD of the mean) in the proximal splice site window (Supplementary Table 1), demonstrating a tendency for all models to predict larger effects for variants close to splice junctions regardless of whether those variants actually disrupt splicing.

### 2.4 sQTL Evaluation

To evaluate the variant effect prediction performance of these models in the endogenous genomic context, we evaluated variant effect prediction using fine-mapped sQTL credible sets from GTEx V8 (49 tissues). Leafcutter [42] and txrevise sQTL fine-mapping results from these tissues were obtained from the eQTL Catalogue [43]. Leafcutter identifies sQTLs from intron excision ratios without reference annotations, while txrevise identifies sQTLs from changes in annotated transcript isoform proportions. We also obtained an independent set of leafcutter sQTL fine-mapping results from human airway epithelial cells (HAEC), the same cell type and dataset used to train SPLAIRE.

To benchmark model performance on a set of presumed causal sQTL variants, fine-mapped variants with posterior inclusion probability (PIP) *>* 0.9 were matched 1:1 to non-causal variants (PIP *<* 0.01) by gene expression (median TPM per tissue) and distance from the splice site (Methods). Variant effect scores were calculated as the maximum per-position absolute difference in predicted splice site probability between reference and alternate sequences within ±2,000 bp of the variant. On the leafcutter credible sets, SpliceAI achieved the highest median AUPRC across 49 tissues (0.79), followed by Pangolin (0.78), SPLAIRE, SPLAIRE-var, and SpliceTransformer (0.76) (*n* = 76–1,815 tested variants per tissue; Figure 6a). On the HAEC dataset, all models had similar performance with AUPRC ranging from 0.69 to 0.72 (*n* = 626; Supplementary Figure S7e). Model rankings were unchanged when the PIP threshold for defining positives was relaxed from 0.9 to 0.5, though absolute AUPRC decreased with the inclusion of lower-confidence positives (Figure 6c).

**Figure 6.**
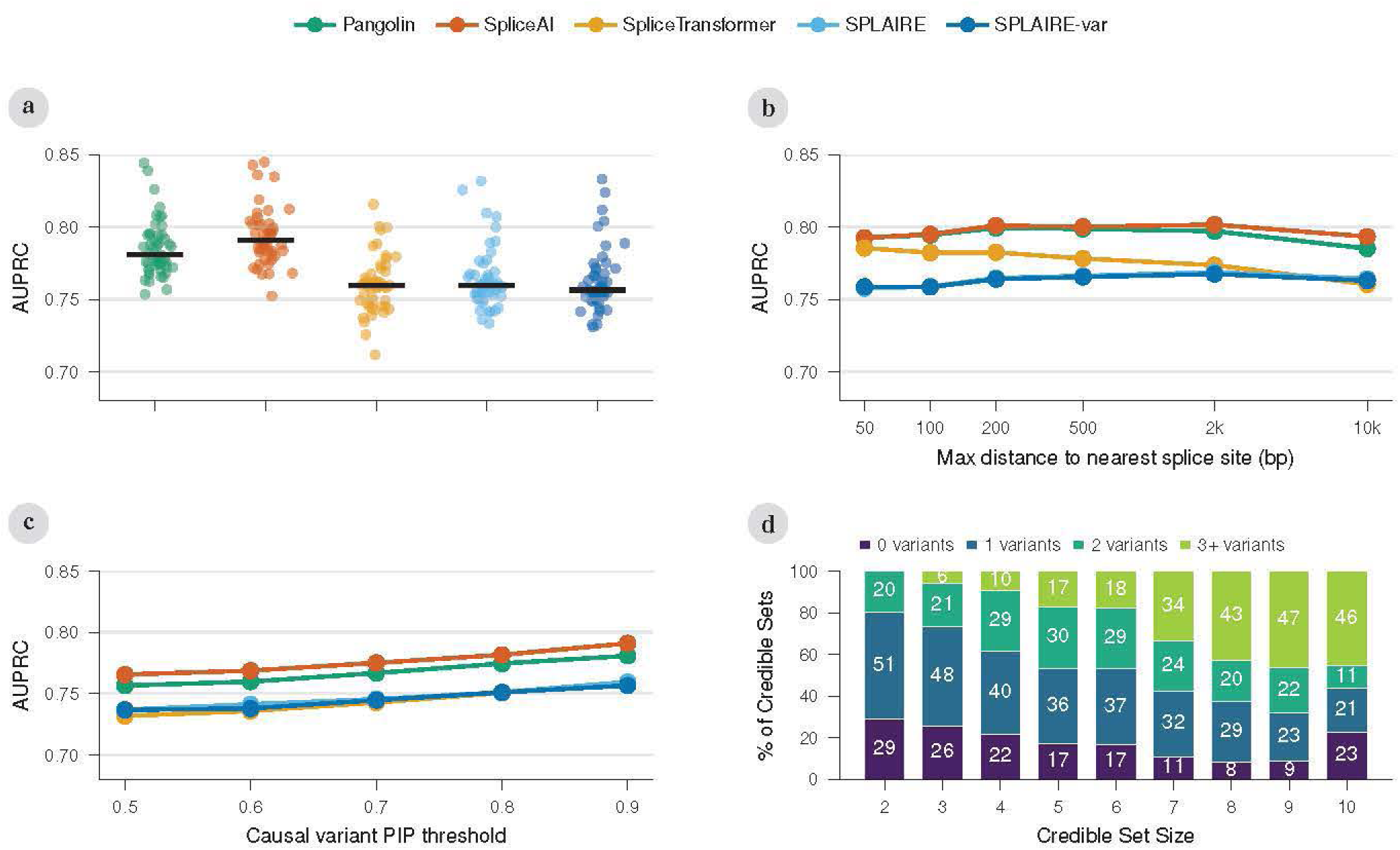
sQTL variant classification. (**a**) Per-tissue AUPRC (points) with median (horizontal line) for classifying fine-mapped leafcutter sQTL causal variants (PIP *>* 0.9) versus matched non-causal variants (PIP *<* 0.01) across 49 GTEx tissues (*n* = 76–1,815 per tissue). (**b**) Cumulative AUPRC stratified by maximum distance of the variant to the nearest splice site. (**c**) Median AUPRC as a function of the PIP threshold used to define positive variants (0.5–0.9). (**d**) Percentage of unresolved credible sets (maximum PIP *<* 0.9, 2–10 variants) in which SPLAIRE calls 0, 1, 2, or 3+ variants positive by credible set size, using per-tissue Youden’s J thresholds [44] derived from the causal versus non-causal classification task (*n* = 7,402 credible sets).

We next examined how model performance varies with distance from the variant to the nearest splice site. For each dataset, we computed AUPRC at cumulative distance thresholds, where each threshold includes all positive-negative pairs at or below that distance. Performance was roughly constant across distance bins for all models on the leafcutter and txrevise datasets (Figure 6b; Supplementary Figure S7b,d,f).

To test whether tissue-specific training improves sQTL detection, we compared the tissue-matched head of each model to its best alternative head within each tissue (SpliceTransformer, 15 tissues; Pangolin, 4 tissues). SpliceTransformer’s tissue-matched head lost to its best alternative head in all 33 tissue comparisons on leafcutter. For the version of Pangolin recommended for variant effect prediction, the tissue-matched head did not outperform alternative heads on leafcutter. The original Pangolin model showed modest tissue-matched gains on leafcutter (6 of 17 comparisons; Supplementary Figure S8). Overall, tissue-matched heads provided little to no advantage over alternative heads for sQTL variant classification.

Performance on txrevise sQTL credible sets was consistent with the leafcutter results (median AUPRC 0.70–0.74; Supplementary Figure S7c). Variants detected as sQTLs by both methods (*n* = 1,498) achieved higher AUPRC than dataset-exclusive variants (Supplementary Figure S9a,b), suggesting these variants have stronger effects on splicing that are detectable regardless of quantification method. However, when the same shared positives were evaluated with leafcutter-matched versus txrevise-matched negatives, all models increased in AUPRC and relative rankings shifted (Supplementary Figure S9c,d), indicating that negative variant selection directly influences both absolute performance and model ranking.

To evaluate how useful these models might be in reducing the number of potentially causal variants within a credible set, we examined the distribution of model scores in unresolved credible sets where fine-mapping did not resolve a single causal variant (maximum PIP *<* 0.9, 2–10 variants per set; *n* = 7,402 total sets). Using per-tissue Youden’s J [44] classification thresholds derived from the causal versus non-causal task, SPLAIRE narrowed credible sets to a single candidate variant in 53% of size-2 sets and 23% of size-10 sets, and to three or fewer candidates in 46% of size-10 sets (Figure 6d). At the more conservative threshold of 0.2, SPLAIRE called at most one variant per set, resolving 22% of size-2 and 32% of size-10 sets to a single candidate (Supplementary Figure S10).

### 2.5 Feature Attribution Analysis

To understand what functional sequence features were used to predict splice sites, we performed attribution analysis on SPLAIRE using DeepLIFT-SHAP on splice sites from the test set of the five tissues (Methods). We stratified splice sites into low (SSU 0–0.1), intermediate (SSU 0.1–0.9), and high (SSU 0.9–1.0) usage sites and computed the mean attribution value at each position relative to acceptor and donor sites (Figure 7a). This showed that both heads yield attribution peaks centered on the splice site dinucleotide motifs, with the classification head having larger attributions in this region while the attributions of the regression head are more dispersed across the input sequence. Additionally, for both heads, low usage sites show lower attribution at splice site motifs and the polypyrimidine tract.

**Figure 7.**
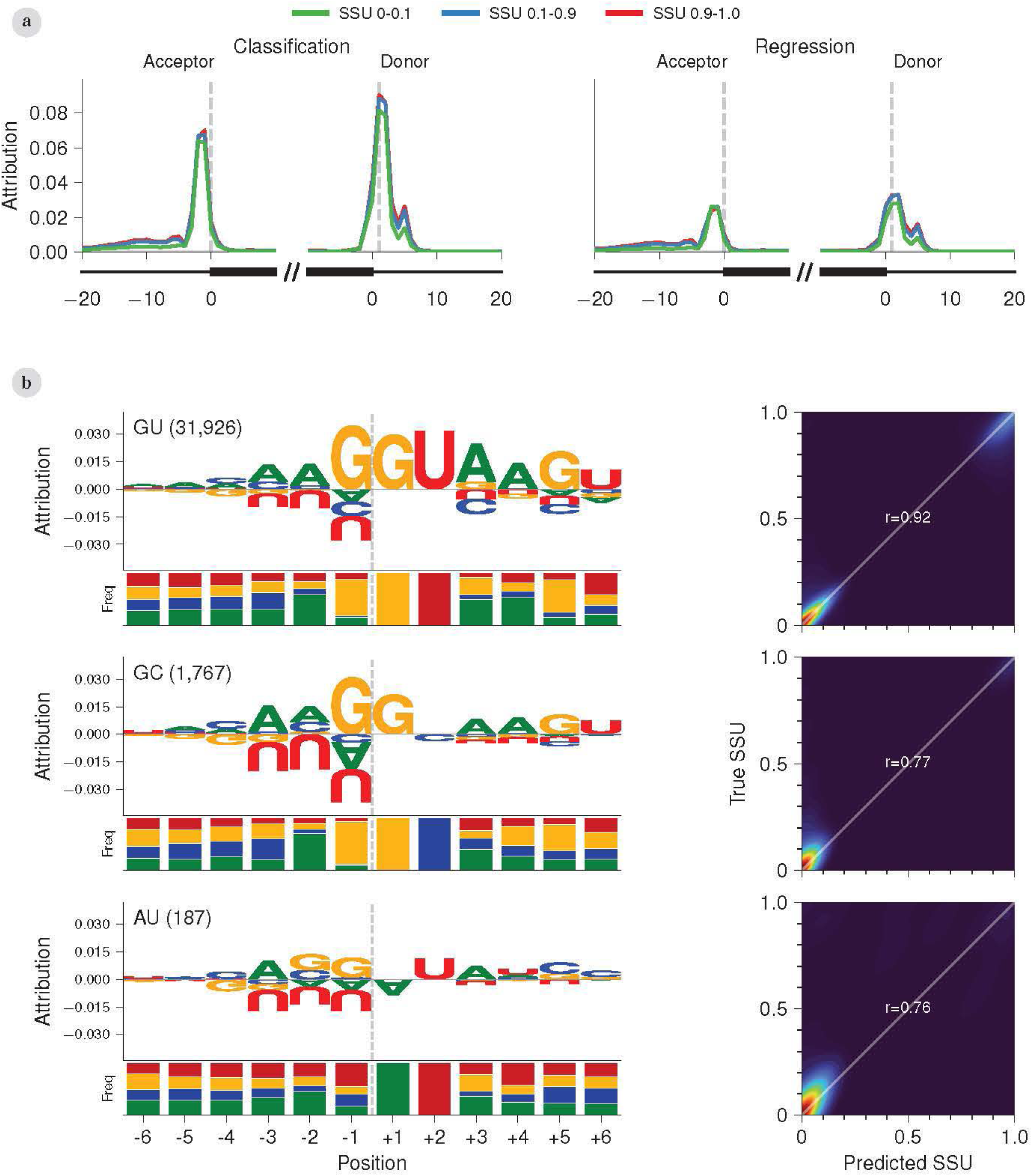
SPLAIRE feature attribution analysis. (**a**) Mean DeepLIFT-SHAP attribution of the classification (left) and regression (right) heads at each position relative to acceptor and donor sites within each SSU group (0.9–1.0 in red, *n* = 10,896; 0.1–0.9 in blue, *n* = 7,709; 0–0.1 in green, *n* = 53,932). (**b**) Donor splice sites grouped by dinucleotide at the first two intronic positions (+1/+2): GU, GC, and AU. For each group, the left column shows the attribution logo from the regression head (top) with a stacked nucleotide frequency bar (bottom) at each position (frequency bar colors match logo letter colors: A green, C blue, G yellow, U red). The right column shows a density plot of predicted versus true SSU with Pearson *r* for that dinucleotide group.

To examine the contribution of canonical and non-canonical splice site motifs to model predictions, we stratified donor and acceptor splice sites by dinucleotide [45, 46] to isolate the minority of splice events that used non-canonical splice site motifs. We then computed the stratified prediction performance and mean regression head attribution value of each nucleotide within a 12 nucleotide window about the splice site. For donor sites, SPLAIRE had good performance in non-canonical splice sites (R2 > 0.76). Examination of the nucleotide-specific attribution scores suggests that for the non-canonical GC sites SPLAIRE’s recognition pattern was essentially the same as for canonical sites. However, for the least common AU sites, the recognition pattern was very different, suggesting that even with relatively few examples the model did learn a distinct pattern of splice site recognition for these noncanonical junctions (Figure 7b). However, the good performance noted for donor motifs did not extend to non-canonical splice acceptors (Supplementary Figure S11), which most likely reflects the higher information content in donor motifs generally.

## 3 Discussion

Neural network-based models that link nucleotide sequence to splicing outcomes have delivered a breakthrough in predictive performance that has already had meaningful impact on medical diagnosis for genetic diseases [7, 14]. However, it is also clear that there is substantial room for improvement in predictive and interpretative accuracy. This paper has a number of informative findings. First, despite excellent top-line performance measures, we show that the performance of state-of-the-art splicing models still has clear performance gaps with respect to low usage and tissue-specific splice sites [47, 48]. Second, our model outperforms all other models on prediction of splice-site usage, likely due to the larger number of observations per splice site afforded by training on 100 individuals from a single cell type, even though other models cover more unique splice sites across species. Third, we report comprehensive tests of leading splicing models for genetic variant interpretation, demonstrating the utility of these models for genetic fine mapping and functional variant identification while also highlighting the need for continued improvement in this area. Finally, we use Shapley-value-based attribution (DeepLIFT-SHAP) to understand which biological signals are effectively captured by our model, demonstrating that these models are capable of capturing subtle biological signals such as usage of non-canonical splice sites.

Given the rapid advancements in deep learning sequence models, developing systematic and unbiased methods of evaluation will catalyze further progress. In the SpliceAI publication [14], the authors report a very high AUPRC and top-K performance, but we demonstrate the dependence of these measures on the composition of splice site usage rates in evaluation datasets. Our paper demonstrates the strong dependence of commonly used performance metrics on splice site usage rates, such that even the best performing models show substantially worse performance when evaluated on low usage sites. This illustrates that splice site prediction is not an entirely solved problem and that application of these models for real world use requires further, task-specific assessments of performance and reliability.

This paper provides one of the most comprehensive evaluations to date of the leading sequence-based splicing models, and this systematic comparison provides some useful insights. First, despite using a more complex transformer-based architecture, SpliceTransformer [21] underperformed the other three CNN-based models in most evaluations. One possible explanation is that the transformer-based architecture used in SpliceTransformer made less efficient use of the available training data than less complex models. Another possibility is that SpliceTransformer was trained to predict a fundamentally different quantity than the Spliser-based SSU used by Pangolin and SPLAIRE. SpliceTransformer defines tissue “usage” for its GTEx training data as the fraction of samples in a tissue containing reads supporting a splice junction [21], a sample-level detection rate that is correlated with but distinct from per-read SSU. Pangolin was generally competitive with SpliceAI and SPLAIRE, though its performance on splice site classification and SSU quantification was often slightly lower. Pangolin’s tissue-specific prediction heads did not show consistent tissue-specific performance as we address later in this section. The original SpliceAI model, which contains only a classification head, offers a useful counterpoint to these multi-task observations. Pangolin and SpliceTransformer use richer training datasets from multiple tissues and species, yet these models often underperformed relative to SpliceAI. The one task where SpliceAI clearly lags is the use of its classification probabilities as direct SSU predictions, which overestimate SSU at weak and intermediate sites [32], an expected limitation given the way that SpliceAI was trained.

The only evaluation that showed one consistently superior model was for SSU quantification, where SPLAIRE outperformed all other models, even in held-out tissues. Since the SPLAIRE model architecture is similar to SpliceAI and Pangolin, the performance boost likely comes from the larger number of training samples available to SPLAIRE (100 distinct subjects), despite the fact that they were all of the same cell type. This suggests that there remains room for improvement in model performance with the use of expanded training sets. We also observed that with Pangolin its regression head provides worse SSU prediction than its classification head. The regression head retains a small rank-correlation advantage (Spearman *ρ* approximately 0.01 higher) but loses on *R*^2^ and MSE, which measure how close predictions are to true SSU values. Both Pangolin and SPLAIRE perform per-head fine-tuning after joint multi-task pretraining [20] (Methods), so the regression head in each model received SSU-specific supervision independent of the classification head. Yet SPLAIRE’s regression head outperforms its classification head while Pangolin’s shows the opposite pattern.

Pangolin and SpliceTransformer are designed to provide tissue-specific predictions, but our analyses demonstrate deficiencies in the tissue-specific performance of these models. Using Pangolin’s test data filtered to tissue-specific sites (1,746 of 36,727), both Pangolin and SpliceTransformer predict target-tissue SSU with lower MAE than SpliceAI and SPLAIRE (Figure 4d; Supplementary Table 10). However, within each model the tissue-matched head is matched or exceeded by its best alternative head on nearly every tissue, indicating that the tissue-specific heads contribute little distinguishing signal beyond what the model already learns at sites with shared usage. To investigate this apparent lack of specificity, we compared DeepLIFT-SHAP attributions across the four Pangolin tissue models at brain-specific splice sites. Attribution patterns are nearly identical across all four models, suggesting that Pangolin’s tissue-specific models tend to identify the same sequence features and assign the same relative importance across tissues. It appears that tissue specificity in Pangolin is primarily encoded in the final 1×1 convolution that maps a shared feature representation to per-tissue SSU. Recent work has explored alternative strategies for learning tissue specificity that include using external information to help models identify tissue-specific sequence elements. PanExonNet [49] proposes conditioning predictions on a learned splicing state derived from RBP and spliceosomal gene expression. TrASPr [50] takes a related approach by conditioning on a learned latent embedding of gene expression and solely predicting a continuous PSI regression target of exons rather than identifying splice sites. Both efforts suggest that tissue-specific splicing prediction likely requires a more informative prior based on expression of splicing factors and careful consideration of training data composition.

One of the most compelling use cases for sequence-based splicing models is for genetic variant interpretation and genetic fine-mapping. Splice-altering genetic variation has been shown to be an important variant class for both Mendelian [51, 52] and complex human diseases [53], and clinical guidelines now incorporate computational splicing predictions into variant interpretation [6, 7]. We observed that for variant effect prediction there was no consistent winner among the models we tested, although when applied to sQTL data all models had reasonable good ability to detect likely causal splicing variants (PIP > 0.9). In fine-mapping contexts where the goal is to narrow the set of potential causal variants, sequence-based splicing models do seem to be useful. The inconsistent model ranking on Vex-seq versus MFASS likely reflects experimental design differences. Vex-seq [41] selected alternatively spliced exons with a sequencing-based ΔPSI readout sensitive to continuous effects, capturing distal effects from splicing enhancers and silencers [54, 55]. MFASS [39] used constitutive exons with a Sort-seq readout designed to detect large-effect splice disruptions, with limited sensitivity to small effects. Both assays use minigene reporters with truncated flanking intronic sequence [39, 41], constraining genomic context. SPLAIRE and SPLAIRE-var significantly outperformed Pangolin and SpliceAI on Vex-seq exonic variants (Figure 5b), while Pangolin achieved the highest correlation and AUPRC on MFASS (Figure 5d). Examining within-model behavior on Vex-seq, classification heads outperformed regression heads on variant effect prediction across most multi-task models. SPLAIRE, SPLAIRE-var, and Pangolin all achieve higher Pearson correlation with measured ΔPSI when scoring with classification heads (SPLAIRE 0.59 versus 0.46, SPLAIRE-var 0.60 versus 0.47, Pangolin 0.55 versus 0.49), while SpliceTransformer is the exception (regression 0.55 versus classification 0.53). On MFASS the differences are uniformly small across multi-task models (within ~0.03 Pearson *r*). The architectural feature most useful for SSU quantification therefore appears poorly aligned with the variant-effect task.

With sequence-based models, one of the main scientific goals is the identification of functional genomic elements and their interaction patterns, which depends on having reliable methods for model interpretation. Our analyses confirmed that the majority of signal is located close to splice sites, consistent with prior observations [5, 56]. However, for these models to achieve the goal of highly accurate prediction of clinically relevant splice variants, they will also need to recognize weaker signals such as non-canonical splice sites and distal functional elements. Our interpretive analysis probed the limits of current models, observing good recognition of non-canonical donor sites but poor recognition of non-canonical acceptor sites. We also observed a substantial dropoff in variant effect detection on reporter assays when considering variants that were not very close to the splice junction. Future advances may require more innovative training approaches that can identify weaker signals that otherwise would be masked by stronger effects around the splice site [30, 57]. Recent alternative training paradigms include event-level cassette-exon transformers focused on adjacent regulatory regions [50] and trans-regulation-aware models that condition predictions on splicing-factor expression states [49], both of which represent promising directions for capturing weaker signals.

Several caveats apply to the interpretation of these results. First, we evaluated three other models, but sequence-based splicing models continue to evolve quickly so this comparative analysis is not comprehensive but relevant mostly to CNN-based models that are closely related to SpliceAI in terms of architecture. We evaluated multiple tissues, but our performance assessment still covers only a small amount of the overall biological range of tissue-specific splicing. DeepLIFT-SHAP requires a baseline distribution, and the choice of baseline directly affects both the magnitude and the ranking of attributions. We use 20 dinucleotide-shuffled baselines per input, which preserves dinucleotide composition and disrupts functional motifs, but other reasonable choices (random sequences, shuffled within-receptive-field windows, or a fixed reference) yield different attribution profiles, and no consensus standard exists in the field. DeepLIFT-Rescale and other gradient-based methods are also known to under-attribute cooperative motifs whose contribution is non-additive [58], a limitation directly relevant to splicing because RNA-binding protein binding events are often cooperative. Univariate Shapley contributions of the type used here cannot capture interactions between distinct binding sites or between motifs and multivariate Shapley methods that quantify such interactions are computationally prohibitive at scale. Finally, our use of L1 normalization to compare attribution magnitudes across SSU bins is a methodological choice rather than an established convention, and absolute attribution comparisons between bins should be interpreted with that caveat in mind.

## 4 Conclusions

Sequence-based deep learning splicing models have advanced computational prediction of splicing and, in turn, the ability to prioritize splice-disruptive variants. Nevertheless our comprehensive analysis reveals that performance gaps remain in several practically important regimes. SPLAIRE, trained on 100 HAEC donors, achieved state-of-the-art SSU quantification across multiple tissues, demonstrating that training methods, specifically data composition, have a significant effect on model performance. Variant effect prediction analysis confounds the importance of classification and SSU regression performance as no single model consistently outperformed on reporter assays ΔPSI prediction or sQTL causality classification. Furthermore, for classification, regression, and variant effect prediction performance we demonstrate that tissue-specific heads in Pangolin and SpliceTransformer provided little advantage over alternative heads. Attribution analysis showed that the models capture canonical and non-canonical splice site motifs and proximal regulatory signals well but place comparatively little weight on distal regulatory elements. Future progress in this field will likely require alternative training objectives, more diverse training data with explicit tissue or cellular context, and approaches that explicitly model cooperative and distal regulatory elements.

## 5 Methods

### 5.1 HAEC-185 cell collection

HAEC-185 is a collection of human primary airway epithelial cells collected from the large airways of 190 individuals during studies investigating regulators of mucus cell hyperplasia. Cells were isolated from human lung tissue obtained primarily from organ donor lungs deemed unsuitable for transplantation due to smoking history, comorbidities, or other clinical factors.

### 5.2 Genotyping and imputation

DNA from all 190 samples was genotyped using the Illumina Global Screening Array. Of these, 183 samples passed quality control and were retained for downstream analyses based on assessments of sex concordance, heterozygosity, genotype missingness, inferred relatedness, and Hardy-Weinberg equilibrium. Genotype data were preprocessed and filtered using *HRC-1000G-check-bim.v4.2.5* (https://www.well.ox.ac.uk/wrayner/tools/) with a reference database derived from the Haplotype Reference Consortium (HRC). Variants that did not match known HRC variants based on genomic position, variant identifier, observed alleles, or minor allele frequency were excluded. Genotype imputation, including liftover to GRCh38, was performed using Minimac4 via the TOPMed Imputation Server (https://imputation.biodatacatalyst.nhlbi.nih.gov/) with the TOPMed r2 reference panel (*apps.topmed-r2@1.0.0; hg38*). Pre-phasing was carried out using Eagle v2.4, with no r2 filter applied (Loh et al., 2016). In total, 307,883,041 variants were imputed.

### 5.3 RNA sequencing

RNA sequencing was performed for 185 HAEC samples. Total RNA was extracted and quantified using the Quant-iT™ RiboGreen® RNA Assay Kit (average RIN=8.4). cDNA libraries were prepared using the Illumina TruSeq™ Stranded mRNA Sample Preparation Kit. Flowcell cluster amplification and sequencing were performed according to the manufacturer’s protocols using the NovaSeq S2 to generate 101bp paired-end reads to an average depth of 50 million mapped reads per sample.

### 5.4 Data preprocessing

Two sets of data were processed for this work. The first was 182 samples of the HAEC-185 dataset. This dataset consists of paired-end short-read RNA-seq coupled with genotyping information for all samples. The second set was 10 samples of lung, testis, whole-blood, and brain-cortex tissue from GTEx v10 release for which there were short-read RNAseq samples paired with genotyping information. The latter is distinct from the dataset used to train the SpliceTransformer models where splice site usage was derived from the exon-exon junction read count file and not processed from short-read RNAseq. Both the HAEC-185 dataset and the GTEx dataset was processed identically. The short-read RNAseq data were aligned to the human reference genome (GRCh38 v45) using STAR v2.7.11b [59]. First-pass alignments were performed for all samples, then a second-pass alignment was performed supplying splice junction files from all samples. Resulting BAM files were filtered to retain only uniquely mapping reads and reads that passed WASP filtering [60] (vW:i:1 tag).

Splice junctions were extracted from these BAM files into BED format using RegTools v1.0.0 [61] “junction extract” command. Each filtered BAM file and corresponding splice junction BED file were then processed with SpliSER v0.1.8 (“process” command) to calculate splice site usage (SSU) for every detected splice site. All splice sites from SpliSER outputs were aggregated across samples to generate a complete list of splice sites. Using a modified version of SpliSER “combine” function, this complete splice site list was used to re-query each BAM file and extract “non-usage” reads at sites present in only a subset of samples. From these combined SpliSER BED files a splice-site-by-sample matrix was constructed in which sites lacking both “usage” and “non-usage” reads were marked as NaN rather than zero SSU.

Using the GENCODE v45 [62] transcriptome annotation, the most upstream transcription start and most downstream transcription end sites for all protein-coding genes were extracted. Overlapping transcripts on the same strand were merged into a single region to prevent overrepresentation during model training. Paralogous genes were identified using the Ensembl BioMart database [63] (GRCh38); genes with at least one human paralog listed were classified as paralogous. For each sample, SSU values and splice site positions were mapped to these transcript regions. Phased variants within each transcript were then extracted from the VCF yielding entries for each transcript-sample-allele combination.

For each remaining transcript-sample-allele entry, the sequence–from transcription start to end–was extracted from the GRCh38 reference genome and reference nucleotides were substituted with alternate nucleotides. Nucleotide sequences were one-hot encoded as follows: A = [1,0,0,0], C = [0,1,0,0], G = [0,0,1,0], T = [0,0,0,1], N (padding) = [0,0,0,0]. Each sequence was zero-padded so its length became a multiple of SL, then further padded with CL/2 bases on both ends (cf. Model architecture for definition of SL and CL). Target arrays of shape SL × 4 were created, where each position consists of [neither, acceptor, donor, SSU]: the first three indices are binary labels for splice sites or other nucleotides, and the fourth index contains the SSU value. Each one-hot encoded nucleotide sequence was split into windows of length CL/2 + SL + CL/2 and the target array is split into windows of length SL. For each window only the central SL positions of the encoded nucleotide sequence map to a target vector of length SL, while the CL/2 bases on both sides serve as contextual information. We also created this same dataset without SNVs inserted in order to train a model on only reference genome sequences and personal sequences.

### 5.5 Model architecture

SPLAIRE and SPLAIRE-var models largely mirror the design of SpliceAI and Pangolin. The model accepts one-hot encoded DNA sequences of length N (where N > CL) and predicts for the central N - CL positions, producing for each base the probability of being an acceptor, donor, or neither, as well as a continuous SSU estimate. The one-hot encoded input is projected to 32 channels by a 1×1 convolution and processed through 16 residual blocks [19] arranged in four groups of four. Successive groups use increasing kernel sizes (11, 11, 21, 41) and dilation rates [17] (1, 4, 10, 25), enabling the network to integrate information from local splice site motifs to distal sequence elements up to 5,000 bp from each predicted position. Each residual block consists of two successive applications of batch normalization, ReLU activation, dropout (rate 0.2), and dilated convolution, with a residual connection. The aggregated skip connection is cropped by CL/2 on each end to yield predictions only for central positions with complete flanking context. The classification head applies a 1×1 convolution with softmax activation, producing per-position probabilities over three classes (acceptor, donor, neither). The regression head applies a 1×1 convolution with a sigmoid activation maps the output to SSU estimates in the [0, 1] interval.

### 5.6 Model training

The training data consisted of pre-mRNA sequences from all autosomal chromosomes (1-22) across 100 individuals from the HAEC cohort. Of the 183 donors that passed quality control, 100 were used for training and 10 for testing (matching GTEx tissue sample counts). All genes (both paralogous and non-paralogous) on chromosomes 2,4,6,8,9-22 were included in training, along with paralogous genes on chromosomes 1,3,5,7. Non-paralogous genes on chromosomes 1,3,5,7 were reserved exclusively for testing. Splice sites annotated in GENCODE v45 but not observed in any RNA-seq sample were added to training targets with masked SSU values. Held-out test data consisted of pre-mRNA sequences from non-paralogous genes on chromosomes 1,3,5,7 across 10 held-out HAEC individuals and 10 individuals from the GTEx v10 cohort across four tissues (Lung, Testis, Whole-blood, and Brain-cortex).

Models were trained with a batch size of 128 using the Adam optimizer [64] with an initial learning rate of 1e-4. An epoch was defined as one tenth of the total training data. The learning rate was reduced by a factor of 0.5 after no improvement in validation loss for 3 epochs, and training was stopped early if validation loss did not improve for 10 epochs. Categorical cross-entropy was used as the loss function for classification predictions and masked binary cross-entropy for regression SSU predictions, where positions with fewer than five supporting reads were excluded from the regression loss. This training was performed using reference sequences for the SPLAIRE model and using sequences with personal SNVs for the SPLAIRE-var model.

After joint training, each model was fine-tuned with only one head and the other frozen, using a reduced learning rate of 1e-5. Fine-tuning used early stopping with a patience of 10 epochs and restored the best model weights. This produced five classification models and five regression models for both SPLAIRE and SPLAIRE-var. All model training was performed on NVIDIA H100 SXM GPUs. All inference for SPLAIRE and comparison models (SpliceAI, Pangolin, SpliceTransformer) was performed on NVIDIA V100-SXM2 GPUs.

### 5.7 Model Evaluation across Tissues

We selected 10 subjects from the GTEx v10 cohort with paired genotyping and RNA-seq data in testis, brain cortex, whole blood, and lung tissues (donor and sample IDs in Supplementary Table 5). The cohort VCF and 40 BAM files were downloaded using the Google Cloud CLI and processed with the same pipeline applied to HAEC BAM files, yielding pre-mRNA sequences with personal genetic variants and per-position splice-site annotations and SSU values. Sequences from non-paralogous protein-coding genes on chromosomes 1, 3, 5, and 7 were used for evaluation. For classification and regression evaluation, we used Pangolin v1. For variant effect prediction, we used Pangolin v2 (fine-tuned on additional data), following the authors’ recommended usage for each task. Throughout this manuscript “Pangolin” refers to the v1 model unless “Pangolin-v2” is stated explicitly.

#### Classification evaluation

AUPRC was computed for each tissue-donor combination, treating annotated splice sites as the positive class and all other nucleotide positions as the negative class. Acceptor and donor splice sites were evaluated together (positives include both site types). Three splice site reference sets were used. Set (i) is all GENCODE v45 splice sites contained in protein-coding transcripts. Set (ii) is the more restrictive set of splice sites in MANE Select transcripts [37]. Set (iii) is all splice sites observed in the RNA-seq data. For each model the classification head was used (SPLAIRE cls head, SpliceAI cls, Pangolin’s per-tissue *p*_splice_ heads averaged, SpliceTransformer’s cls channel). Per-tissue and per-donor classification metrics for all individual heads are tabulated in Supplementary Tables 6 and 8. AUPRC was additionally stratified by SSU decile. Splice sites were binned into ten equal-width bins from 0 to 1 and AUPRC was recomputed within each bin (Figure 3b,d).

#### Regression evaluation

For each splice site with a SSU measurement (at least five supporting reads in the donor sample), we computed Pearson correlation, Spearman correlation, *R*^2^, MSE, and MAE between predicted and measured SSU. Acceptor and donor sites were treated as separate observations. SPLAIRE’s regression head produces a continuous SSU prediction directly. SpliceAI does not have a regression head, thus we used the maximum of its acceptor and donor classification probabilities as an SSU proxy. Pangolin and SpliceTransformer each have multiple tissue-specific regression heads. Figure 3e–h shows Pangolin’s four tissue heads averaged and SpliceTransformer’s 15 tissue heads averaged. Metrics for all individual heads are tabulated across four SSU subsets. The four subsets are progressively more restrictive. *Valid* sites have a measurable SSU in the donor sample. *Valid_>_*_0_ is the valid subset restricted to sites with measured SSU *>* 0. Per-tissue per-head metrics on *Valid* and *Valid_>_*_0_ sites are tabulated side-by-side in Supplementary Table 7; the corresponding *Shared* and *Shared_>_*_0_ tables are in the additional supplements. MSE was additionally stratified by SSU decile (Figure 3f,h).

#### Annotated versus unannotated subset

Splice sites observed in the RNA-seq data were partitioned into *annotated* (also present in the GENCODE v45 primary assembly annotation) and *unannotated* subsets, and regression metrics were computed within each subset (Supplementary Table 4; Supplementary Figure S4).

#### Pangolin’s published test set

To evaluate models on independent test data, we additionally scored all models on the four-tissue (heart, liver, brain, testis) test set published with Pangolin [20]. The test data were obtained from the Pangolin training repository (https://github.com/tkzeng/Pangolin_train) and consist of splice sites in non-paralogous human protein-coding genes on chromosomes 1, 3, 5, 7, and 9, with SSU values quantified by SpliSER from the RNA-seq data of Cardoso-Moreira et al. [34]. We restricted evaluation to chromosomes 1, 3, 5, and 7 to match SPLAIRE’s held-out test chromosomes. We additionally applied our paralog filter to Pangolin’s test set, removing all genes with at least one human paralog under our paralog annotation. Results both with and without this additional filter are reported (Supplementary Figure S3; Supplementary Table 9).

#### Calibration analysis

To assess model calibration on continuous SSU prediction, we plotted measured versus predicted SSU per model and tissue and fit both a LOESS curve and a linear regression. Linear-fit slope and intercept summarize systematic over- or under-prediction across the SSU range, with slope of 1 and intercept of 0 indicating perfect calibration (Supplementary Figure S2; Supplementary Table 2).

#### Tissue specificity analysis

We applied the *τ* tissue-specificity index [38] to splice sites with measurable SSU across multiple tissues:

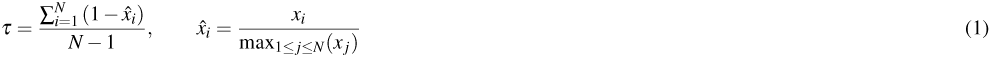

where *x_i_* is the mean SSU in tissue *i* (averaged across held-out individuals) and *N* is the number of tissues in the analysis. *τ* ranges from 0 (constitutive) to 1 (perfectly tissue-specific). We additionally computed a reverse variant *τ*_rev_ by evaluating the same formula on the inverted values (1 − *x_i_*) to capture exclusion-specific sites where one tissue shows unusually low SSU while others are high. Sites were classified into three groups: constitutive (*τ <* 0.1), intermediate (0.1 *τ ≤* 0.5 and *τ*_rev_ *<* 0.7), and tissue-specific (*τ* ≥ 0.5 or *τ*_rev_ ≥ 0.7, with the additional requirement max*_i_*(*x_i_*) − min*_i_*(*x_i_*) ≥ 0.3). Sites with *τ <* 0.1 and elevated *τ*_rev_ were retained in the constitutive group as they have range *<* 0.3.

For each tissue-specific site we identified the *target tissue* as the tissue whose mean SSU deviated most from the cross-tissue mean, and classified the site as inclusion-specific (higher SSU) or exclusion-specific (lower SSU). We then computed per-target-tissue MAE between predicted and measured SSU within each specificity group. For models with multiple tissue-specific heads (Pangolin, SpliceTransformer), we evaluated both the *tissue-matched* head (the head corresponding to the target tissue) and the *best alternative* head (the non-matched head with lowest MAE for that target tissue). SpliceTransformer is omitted for testis as it does not have a testis-specific head. SPLAIRE and SpliceAI have only tissue-independent heads and are not included in this matched-vs-alternative comparison.

This analysis was performed on three datasets. Figure 4 uses Pangolin’s published 4-tissue test set (heart, liver, brain, testis, *N* = 4, 36,727 shared sites) before our additional paralog filter, matching the data on which Pangolin’s authors evaluated their own model. The same analysis was repeated on our 5-tissue evaluation cohort (HAEC, lung, testis, brain cortex, whole blood, *N* = 5, splice sites detected in all five tissues), with results reported in Supplementary Table 10.

### 5.8 Reporter Assays

The reporter assays measure exon-inclusion ΔPSI, the change in the fraction of transcripts that include the tested exon, distinct from SSU. Vex-seq data (2,055 variants with measured ΔPSI) was obtained from the MMSplice repository [41]. The data originates from the CAGI5 Vex-seq challenge [65] and includes both SNVs and indels across 110 alternatively spliced exons assayed in HepG2 cells. MFASS data (27,733 variants with measured ΔPSI) was obtained from the MFASS repository [39] and consists of ExAC SNVs across 2,198 constitutive exons in HEK293T cells.

For each variant, we extracted sequences of length 10,001 bp from the hg19 reference genome centered on the exon start and end with the reference or alternate allele. Sequences on the reverse strand were reverse complemented, one-hot encoded, and scored with SpliceAI, Pangolin, SpliceTransformer, and SPLAIRE.

For each variant we scored the reference and alternate sequences with each model. The per-variant predicted ΔPSI was computed at the acceptor and donor sites of the tested exon as the difference between alternate and reference scores, averaged across the two splice sites. The output channels used per model were as follows. For SpliceAI we used the acceptor and donor classification probability channels. For SPLAIRE and SPLAIRE-var we used the acceptor and donor channels of the classification head. For SpliceTransformer we used the cross-tissue acceptor and donor classification channels. For Pangolin and Pangolin-v2 we used the four tissue-specific *p*_splice_ classification heads (brain, heart, liver, testis) and took the maximum absolute delta across tissues. Model performance was assessed by comparing measured and predicted ΔPSI using Pearson correlation, Spearman correlation, and AUPRC. Pearson and Spearman were computed on all variants within each variant-location category. AUPRC was computed treating splice-disrupting variants (ΔPSI *>* 0.10 for Vex-seq, *>* 0.50 for MFASS) as positives and remaining variants as negatives.

Variants were stratified by location relative to the nearest splice site of the given exon using the categories defined by Mount et al. [65]. Splice site proximal variants were defined as those within 3 nucleotides of the exonic side or 8 nucleotides of the intronic side of each splice junction. Remaining variants were classified as exonic if located within the exon or intronic otherwise. All metrics were computed within each location category.

Statistical significance of pairwise model differences was assessed using paired bootstrap resampling (10,000 iterations). In each iteration, variants were resampled with replacement and all four metrics (Pearson correlation, Spearman correlation, *R*^2^, and AUPRC) were computed for every model on the same resampled set. For each model pair, the bootstrap distribution of metric differences was used to compute two-sided *p*-values and 95% percentile confidence intervals. *P*-values were corrected across 4 comparisons (each model versus the best-performing model) using the Holm–Bonferroni method.

### 5.9 sQTL

Fine-mapped txrevise and leafcutter [42] sQTL credible sets and association summary statistics for 49 tissues from GTEx V8 were obtained from the eQTL Catalogue (https://www.ebi.ac.uk/eqtl/). Gene expression association summary statistics for all tissues were also obtained along with the GENCODE v39 genome annotation. HAEC leafcutter sQTL credible sets and summary statistics were generated from the HAEC-185 RNA-seq and imputed genotype data described above. Splice junction usage was quantified using leafcutter [42] with the GENCODE v45 annotation, intron excision ratios were tested for association with genotype using tensorQTL [66] under an additive linear model, and credible sets were identified by fine-mapping with SuSiE [67].

Gene expression levels were obtained from GTEx V8 median TPM values per tissue. For each tissue, genes were assigned to expression bins by computing log_2_(TPM) and discretizing into bins of width 0.4, such that genes with similar expression levels (within ~1.3-fold) were grouped together.

For each sQTL dataset, a set of causal (positive) and distance and expression matched non-causal (negative) SNVs were constructed to evaluate model performance. Positive variants were selected from fine-mapped credible set variants with PIP ≥ 0.9. For txrevise, positive variants were further required to have “contained” in the molecular trait ID. For all variants, splice distance was defined as the minimum distance from the variant position to the nearest exon boundary within the same gene (for txrevise) or to either boundary of the associated intron (for leafcutter). All variants (positive and negative) were required to have splice distance ≤ 10 kb.

Each positive variant was matched to one negative variant per tissue, selected without replacement. For the GTEx datasets, negative candidates were drawn from gene expression association summary statistics and for HAEC negatives were drawn from the sQTL summary statistics. Negatives were further restricted to SNVs with PIP *<* 0.01. Within each expression bin, positives and negatives were paired one-to-one to minimize total log-distance mismatch using scipy.optimize.linear_sum_assignment. This produces matched pairs with standardized mean differences in splice distance below 0.03 across all three datasets.

For each variant, we extracted 2,000 bp upstream and downstream of the variant position from the GRCh38 reference genome and reverse complemented if the phenotype was on the antisense strand. Sequences were one-hot encoded and scored using SpliceAI, Pangolin (v1 and v2), SpliceTransformer, SPLAIRE and SPLAIRE-var. Reference scores were subtracted from alternate scores and the predicted effect was computed within the ±2,000 bp window. For SpliceAI, the score was the maximum absolute delta across the three classification channels (no-splice, acceptor, donor). For SPLAIRE and SPLAIRE-var, the score was the maximum absolute delta across the three classification channels (no-splice, acceptor, donor). For Pangolin-v2 (used in main figures), the score was the maximum absolute delta across the four tissue-specific *p*_splice_ classification heads (brain, heart, liver, testis). For SpliceTransformer, the score was the maximum absolute delta across all output channels. For each model we additionally computed alternative delta-score formulations using other heads (SPLAIRE SSU regression, Pangolin usage heads, SpliceTransformer per-tissue channels). Model performance was assessed using area under the precision-recall curve (AUPRC) for each tissue. Results for all delta-score formulations are shown in Supplementary Table 3. To assess the performance of models as a function of distance to splice sites, we computed AUPRC at cumulative splice distance thresholds (≤50, ≤100, ≤200, ≤500, ≤2,000, and ≤10,000 bp), where each threshold includes all positive-negative pairs with positive variant splice distance at or below that value.

To test whether the tissue-matched head of each multi-tissue model improves classification on the matching GTEx tissue, we computed per-tissue AUPRC for every individual head and compared the tissue-matched head against the best alternative head from the same model. SpliceTransformer’s 15 tissue heads were mapped to GTEx tissues by tissue name, with GTEx tissues sharing a broader SMTS category (e.g. the 13 SMTSD brain subregions all mapping to the SpliceTransformer brain head) sharing the corresponding tissue head. The four tissue heads of Pangolin-v2 (brain, heart, liver, testis) were mapped to GTEx tissues in the same manner. GTEx tissues without a corresponding tissue head were excluded. Aggregate (max-across-tissues) outputs were excluded from the alternative pool.

To test how sensitive sQTL classification performance is to the choice of positive and negative variant sets, we compared classification on shared versus dataset-exclusive positives across the GTEx leafcutter and txrevise datasets. Positive variants from each dataset were matched by genomic coordinates and reference and alternate bases to partition into a shared set and two dataset-exclusive sets.

To assess model performance in unresolved credible sets, we used GTEx leafcutter credible sets where the maximum PIP was less than 0.9, all variants were SNVs, all variants were within 5 kb of both splice sites of the intron phenotype, and the credible set contained between 2 and 10 variants. Variants in each credible set were scored as described above. SPLAIRE scores were thresholded at the median per-tissue Youden’s J value from the GTEx leafcutter sQTL classification task (0.031 for SPLAIRE) and at a threshold of 0.2. For each credible set we counted the number of variants whose SPLAIRE score exceeded the threshold, and computed the fraction of credible sets with 0, 1, 2, or 3+ variants exceeding the threshold, stratified by credible-set size.

### 5.10 Attribution Analysis

We computed per-nucleotide attributions using DeepLIFT-SHAP [58,68], which measures each input base’s contribution relative to a set of reference sequences. Splice sites from test chromosomes (chr1, chr3, chr5, chr7) in non-paralogous protein-coding genes were selected for this analysis. Sites with ambiguous (“N”) bases in the surrounding genomic sequence were excluded, resulting in 38,456 acceptor and 34,081 donor splice sites. For each site, we extracted a 10,001 bp window centered on the splice site and one-hot encoded the sequence.

DeepLIFT-SHAP requires a set of reference (baseline) sequences against which each input is compared. We generated 20 reference sequences per input by dinucleotide shuffling. Attributions were averaged across all 20 references following the SHAP framework. Attributions were computed for both the regression and classification heads of the SPLAIRE model using the tangermeme library v1.0.0. The model was converted to PyTorch [69] and scored with batch sizes of 64 sequences on NVIDIA V100-SXM2 GPUs.

The attribution method produces scores for all four possible nucleotides at each position (hypothetical attributions), reflecting what each base would contribute if it were present. To obtain the actual contribution of each position, we multiplied the hypothetical attributions element-wise by the one-hot encoded input sequence and summed across nucleotide channels, yielding a single attribution value per position (observed attributions). To enable comparison across sequences with different prediction magnitudes, we applied L1 normalization by dividing each position’s attribution by the sum of absolute attribution values across the 10,001 bp window.

Per-position mean attribution profiles (Figure 7a) were computed as the mean of L1-normalized observed attribution across sites within each SSU stratum, separately for acceptor and donor sites and for the classification and regression heads. Splice sites were stratified into three SSU bins (0.0–0.1, 0.1–0.9, 0.9–1.0).

For the dinucleotide-stratified attribution analysis (Figure 7b and Supplementary Figure S11), donor splice sites were grouped by the dinucleotide at the first two intronic positions (+1/+2, GU, GC, AU) and acceptor sites were grouped by the dinucleotide at the last two intronic positions (2/ 1, AG, AC). Within each group, attribution logos were generated by stacking observed-attribution values per nucleotide at each position, and a stacked nucleotide-frequency bar was computed per position. Per-group Pearson correlation between predicted and true SSU was computed across all sites in the group.

Two attribution metrics were computed per site (Supplementary Figure S12). *Effective width* is the minimum number of positions whose absolute attribution captures 90% of total absolute attribution across the 10,001 bp window. *Positive attribution fraction* is the fraction of total absolute attribution contributed by positions with positive observed attribution. Both metrics were computed independently for the classification and regression heads, stratified by splice site type (acceptor, donor) and SSU bin (0.0–0.1, 0.1–0.9, 0.9–1.0). Pairwise comparisons between SSU bins within each splice-site type used Mann–Whitney *U* tests, with *p*-values corrected by the Holm–Bonferroni method [70].

We additionally computed DeepLIFT-SHAP attributions for each of Pangolin’s four tissue-specific usage heads (brain, heart, liver, testis) at the tissue-specific splice sites defined above (see *Tissue specificity analysis*). The attribution computation followed the same pipeline as for SPLAIRE, using 20 dinucleotide-shuffled reference sequences per input, 10,001 bp windows centered on each splice site, and observed attributions computed by element-wise multiplication with the one-hot input. SPLAIRE-regression attributions were also computed on the same set of tissue-specific splice sites for comparison with Pangolin.

Attribution similarity across models was quantified by computing pairwise Pearson correlation between the per-position attribution vectors of two models at each site. We used both raw observed attributions and L1-normalized attributions, and computed correlation in two windows: the full 10,001 bp input sequence and a ±50 bp window centered on the splice site (Supplementary Figure S14). For each tissue-specific site, the per-pair Pearson *r* values were aggregated by the site’s target tissue (the tissue whose mean SSU deviated most from the cross-tissue mean), producing one violin per (target-tissue, model-pair) combination. We performed two complementary comparisons. First, we compared Pangolin’s four tissue-specific heads against each other (Supplementary Figures S6, S14). Second, we compared SPLAIRE’s regression head against each Pangolin tissue head matched to the site’s target tissue (Supplementary Figure S13).

## Declarations

### Ethics approval and consent to participate

Human airway epithelial cells were derived from lung tissue obtained from deceased organ donors whose lungs were deemed unsuitable for transplantation. [PENDING. add the IRB protocol number and approving ethics committee, or a statement that the study was determined not to constitute human subjects research]

### Consent for publication

Not applicable.

### Availability of data and materials

Processed data, model weights, and predictions are available on Zenodo (https://doi.org/10.5281/zenodo.191364 GTEx v10 data are available through the GTEx Portal (https://gtexportal.org) and dbGaP (phs000424). Vex-seq reporter assay data were obtained from [41]. MFASS reporter assay data were obtained from [39]. Fine-mapped sQTL credible sets and summary statistics were obtained from the eQTL Catalogue (https://www.ebi.ac.uk/eqtl/). Source code is available at https://github.com/NNeuralDynamics/splaire with an archived version on Zenodo (https://doi.org/10.5281/zenodo.19136478) and is released under the MIT License.

### Authors’ contributions

Conceptualization was by M.R., S.G., P.J.C., and A.P. Methodology was by M.R., J.D., P.R., A.P., and S.G. SPLAIRE model development and implementation, software, formal analysis, validation, and visualization were by M.R. Investigation was by M.R., Y.T., D.G.-M., A.S., and C.L. Data curation was by M.R., D.G.-M., A.S., and C.L. Resources were provided by Y.T. Supervision was by J.D., P.R., P.J.C., and A.P. Project administration was by A.P. Funding acquisition was by Y.T., P.J.C., and A.P. The initial draft was written by M.R., P.J.C., and A.P. All authors critically reviewed and edited the manuscript.

## Acknowledgements

Not applicable.

---

**Figure S1.**
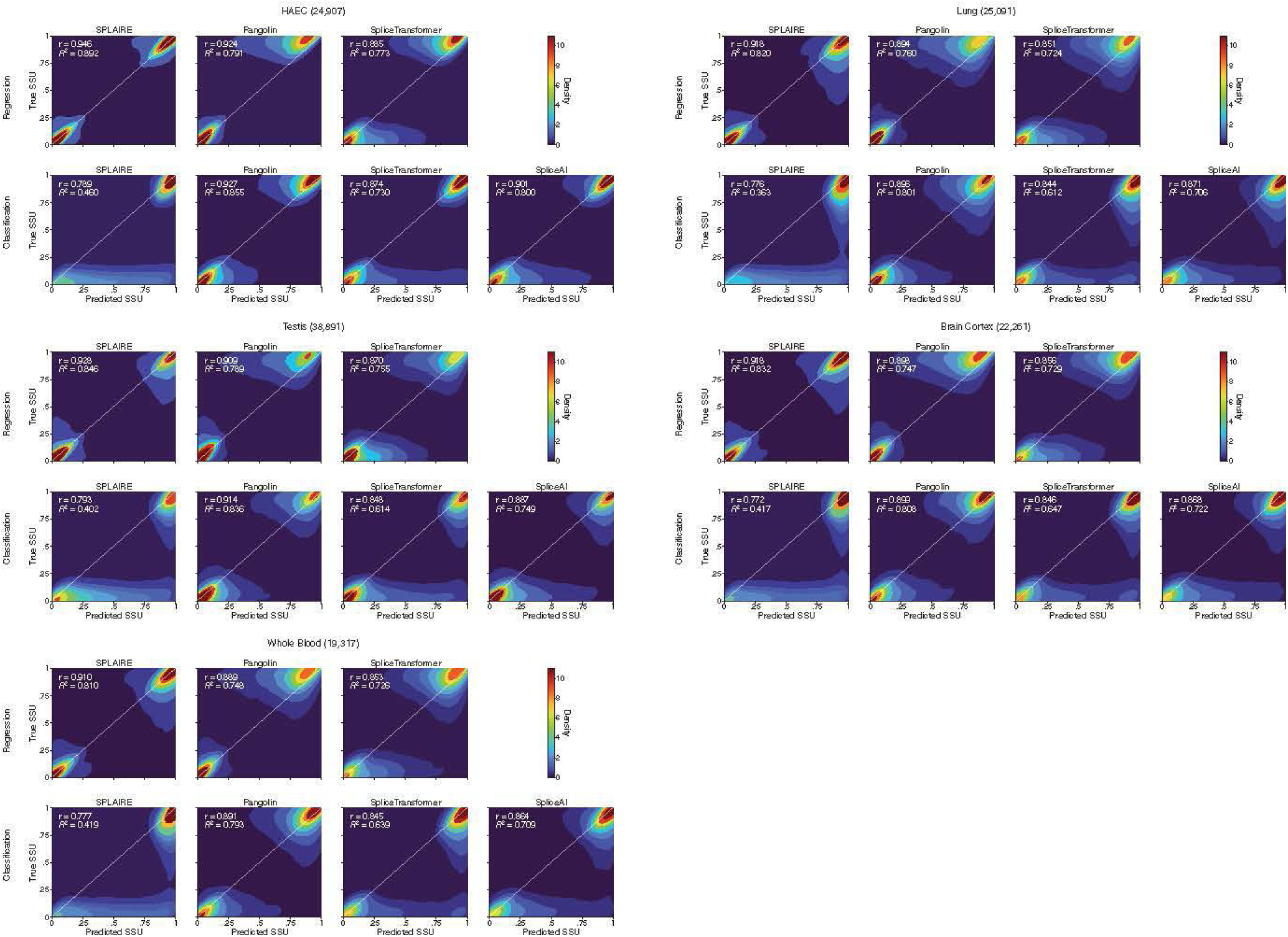
Predicted vs measured SSU density plots by model and tissue. KDE density plots of predicted SSU (x-axis) against measured SSU (y-axis) for all models across held-out individuals. Within each tissue sub-panel, the top row shows regression outputs and the bottom row shows classification outputs. The Pangolin usage and classification outputs and the SpliceTransformer regression output are averaged across tissues. SPLAIRE, SpliceTransformer, and SpliceAI classification outputs are the maximum of acceptor and donor probabilities. Pearson *r* and *R*^2^ are shown per panel.

**Figure S2.**
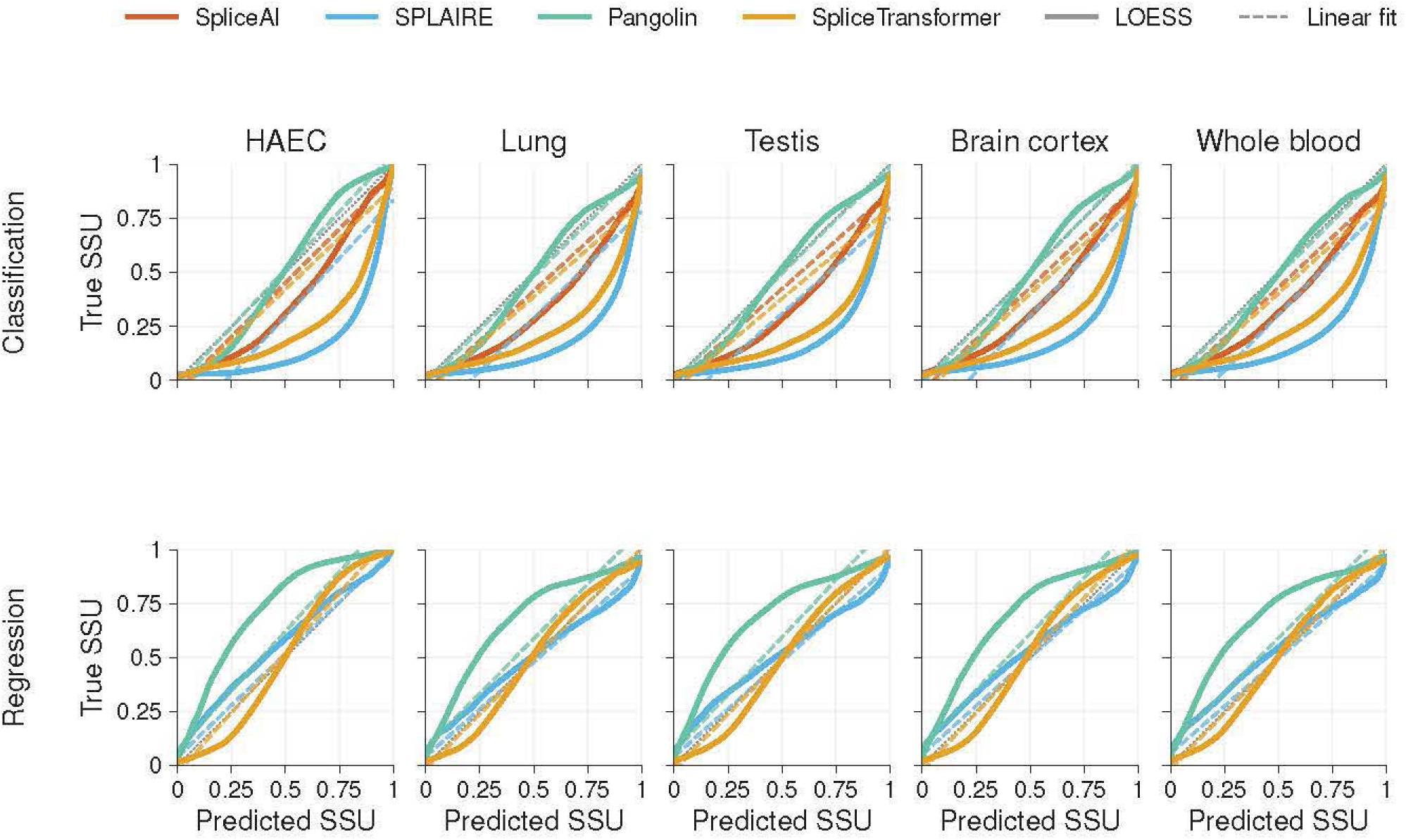
LOESS calibration curves for predicted SSU. For each model and tissue, a LOESS curve (solid) of true SSU (y-axis) against predicted SSU (x-axis) is shown alongside the OLS linear fit (dashed, same color); the black dotted line is the identity. The top row shows classification outputs and the bottom row shows regression outputs. The Pangolin usage and classification outputs and the SpliceTransformer regression output are averaged across tissues. SPLAIRE, SpliceTransformer, and SpliceAI classification outputs are the maximum of acceptor and donor probabilities. Computed on all splice sites (same subset as Supplementary Figure S1), averaged across held-out individuals.

**Figure S3.**
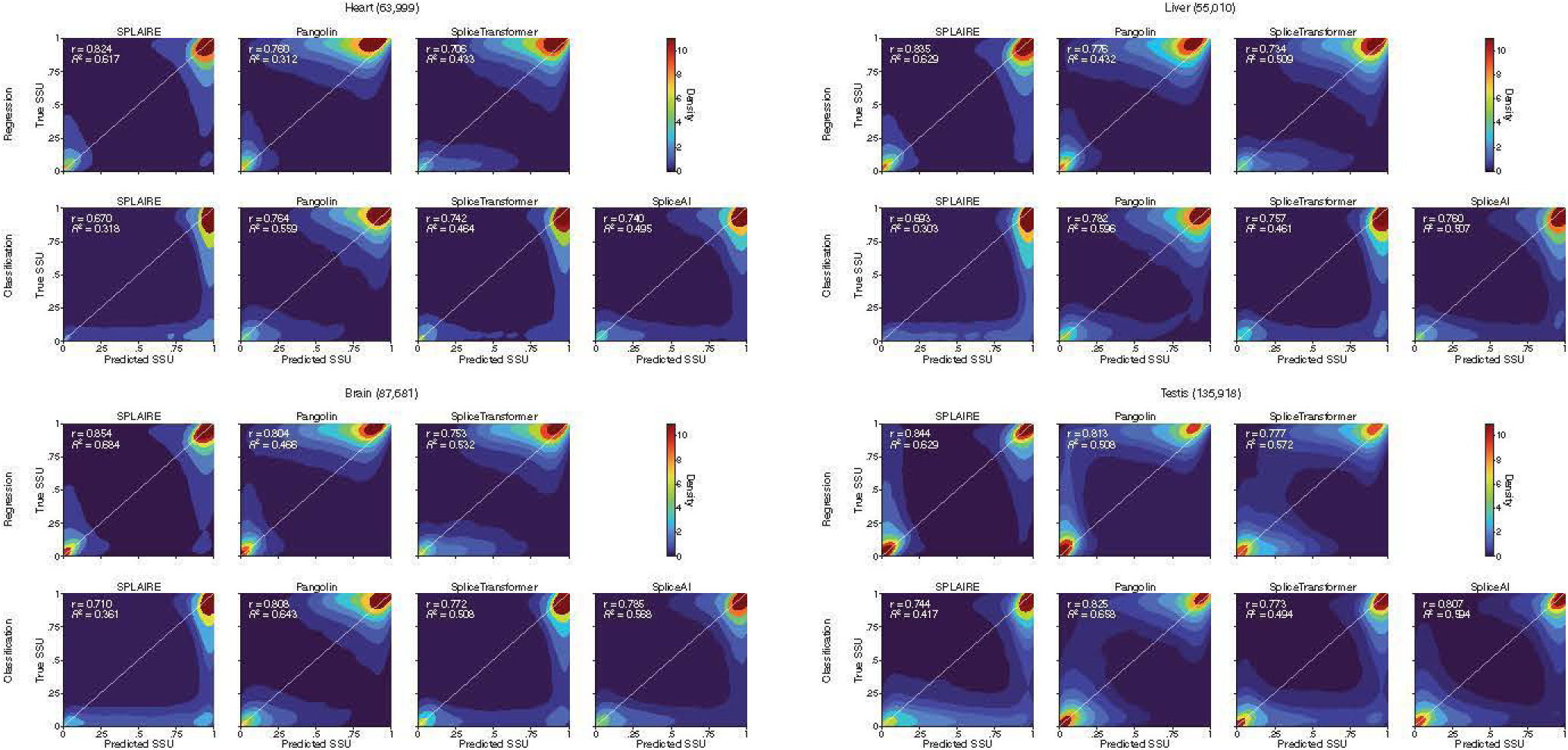
Predicted vs measured SSU density plots on the Pangolin test set. KDE density plots of predicted SSU versus measured SSU on the Pangolin test set. Within each tissue, the top row shows regression outputs and the bottom row shows classification outputs (same layout as Supplementary Figure S1). The Pangolin usage and classification outputs and the SpliceTransformer regression output are averaged across tissues. SPLAIRE, SpliceTransformer, and SpliceAI classification outputs are the maximum of acceptor and donor probabilities. Pearson *r* and *R*^2^ are shown per panel.

**Figure S4.**
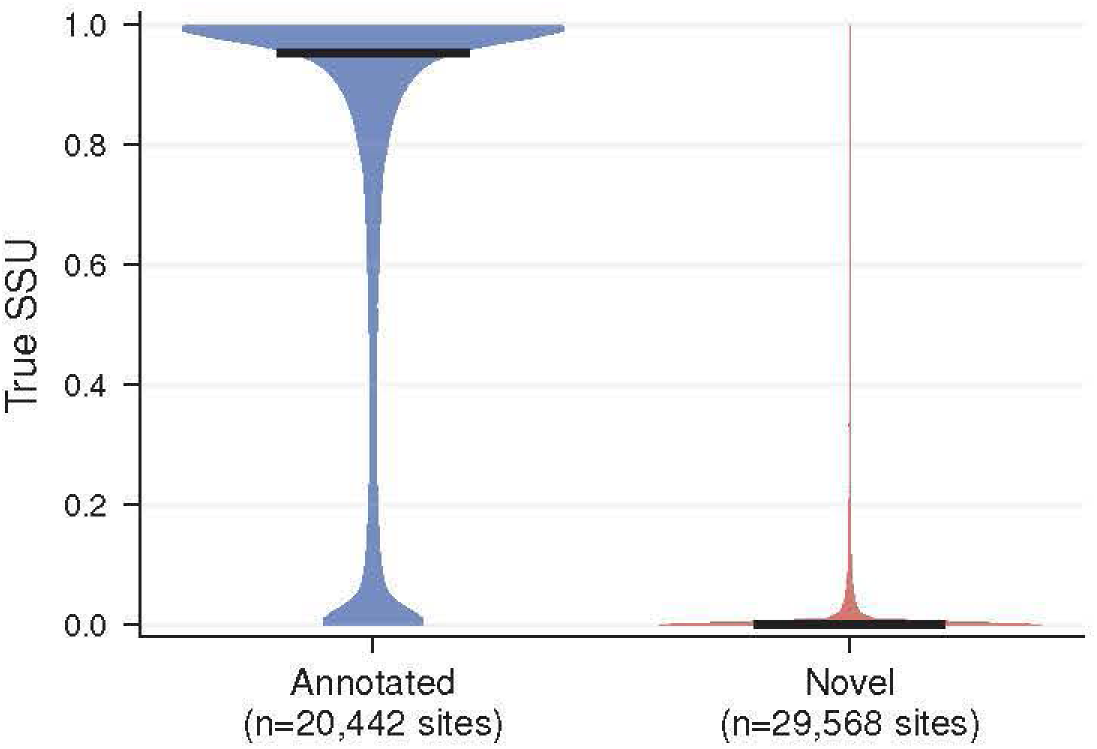
Measured SSU distribution by annotation status across tissues. Violin plot of measured SSU stratified by annotation status (annotated vs unannotated) across HAEC and four GTEx tissues. Annotated sites are also present in the GENCODE v45 primary assembly annotation; unannotated sites are observed in RNA-seq but absent from the annotation.

**Figure S5.**
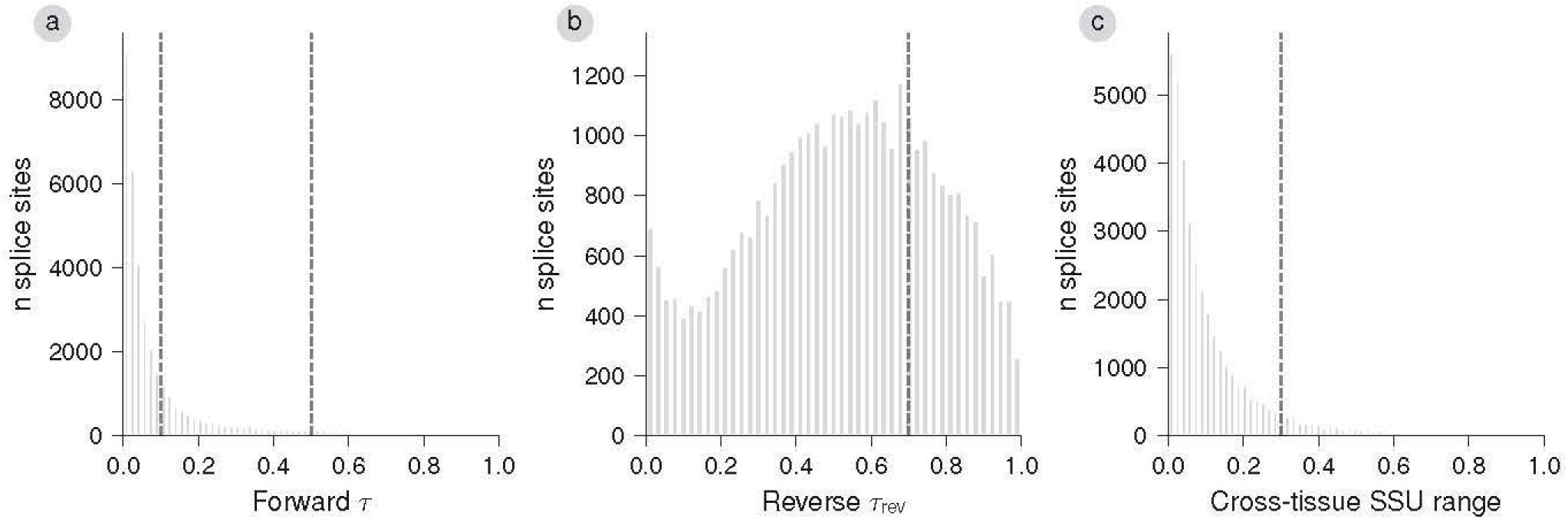
Tissue specificity index distributions on the Pangolin test set. Distributions of forward *τ*, reverse *τ*_rev_, and cross-tissue SSU range used to define the constitutive, intermediate, and tissue-specific groups in Figure 4. (**a**) Forward *τ* distribution with group thresholds at 0.1 and 0.5 (dashed lines). (**b**) Reverse *τ*_rev_ distribution with threshold at 0.7. (**c**) Cross-tissue SSU range (max min SSU across tissues) distribution with threshold at 0.3.

**Figure S6.**
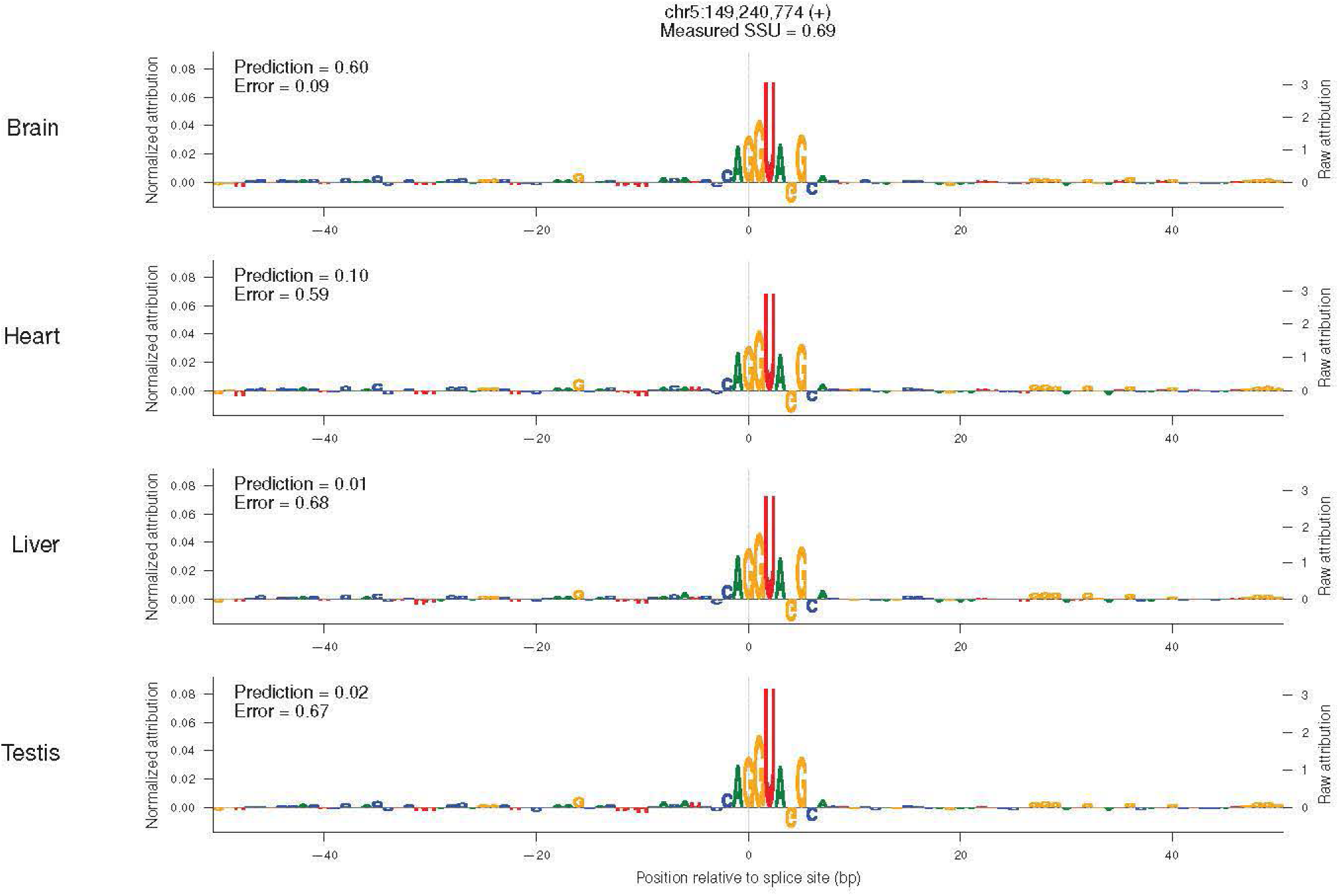
Pangolin attributions at brain-specific splice sites. DeepLIFT-SHAP attribution logos from all four independently trained Pangolin tissue-specific usage models (brain, heart, liver, testis) at a representative brain-specific splice site. Genomic coordinates and measured brain SSU are shown in the title. Each row shows one model’s attributions centered on the splice site (dashed line). Logo letter heights are L1-normalized attributions (left y-axis); the right y-axis shows the corresponding raw (unnormalized) attribution scale for each panel. Per-model predicted SSU and absolute prediction error against the measured brain SSU are annotated in each panel.

**Figure S7.**
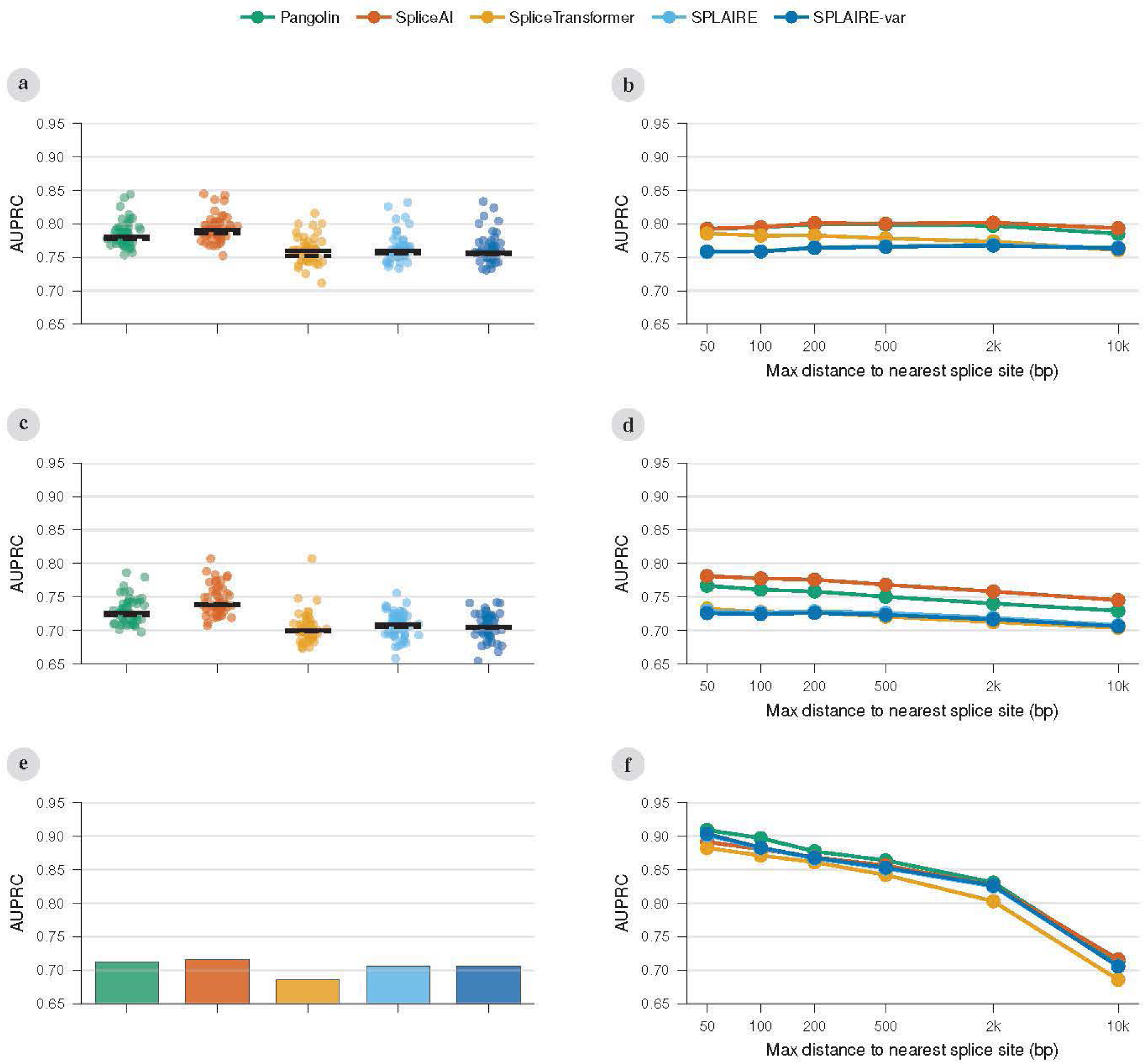
sQTL variant classification across additional benchmarks and metrics. All panels share the same legend (top) as Figure 6. (**a**) Per-tissue AUPRC (points) with median (black bar) for classifying fine-mapped leafcutter sQTL causal variants versus matched non-causal variants across 49 GTEx tissues. (**b**) Cumulative AUPRC by maximum splice distance of the causal variant to the nearest intron boundary for GTEx leafcutter. Panels (a) and (b) present the data from Figure 6a,b. (**c**) Per-tissue AUPRC for classifying causal versus non-causal variants from GTEx txrevise sQTL credible sets (*n* = 63–1,054 per tissue). (**d**) Cumulative AUPRC by splice distance for GTEx txrevise. (e) AUPRC for classifying causal versus non-causal variants from a HAEC leafcutter sQTL dataset (*n* = 626). Single tissue, shown as bars. (f) Cumulative AUPRC by splice distance for HAEC.

**Figure S8.**
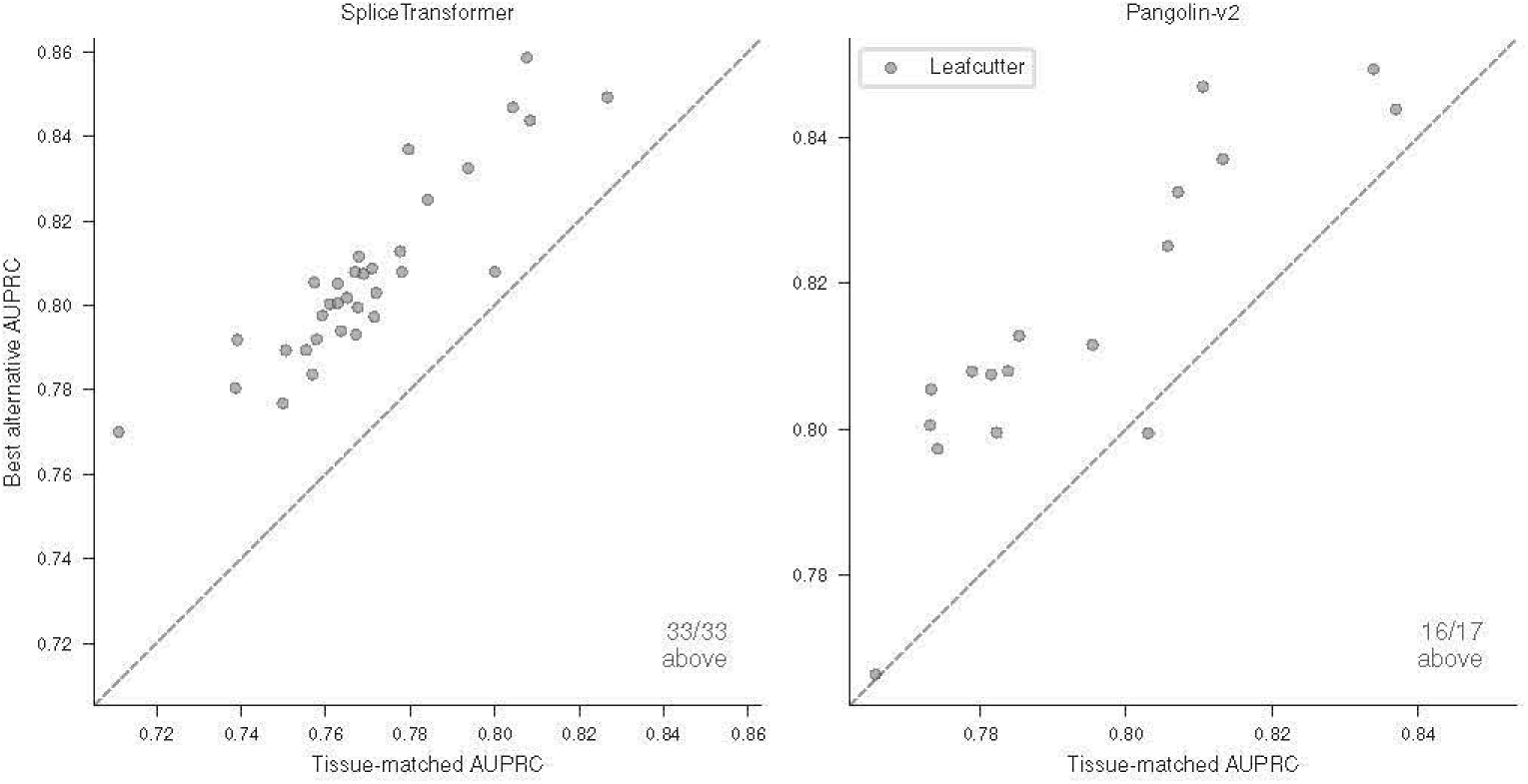
sQTL classification AUPRC by tissue-specific output head. Tissue-matched head AUPRC (x-axis) versus best alternative head AUPRC (y-axis) for SpliceTransformer (15 tissue heads), Pangolin (4 tissue heads), and Pangolin v2 (4 tissue heads) on leafcutter sQTL credible sets. Each point is one GTEx tissue. Points above the diagonal indicate the tissue-matched head is outperformed by another individual tissue head. Aggregate (max-across-tissues) outputs are excluded. Counts above diagonal are annotated per panel.

**Figure S9.**
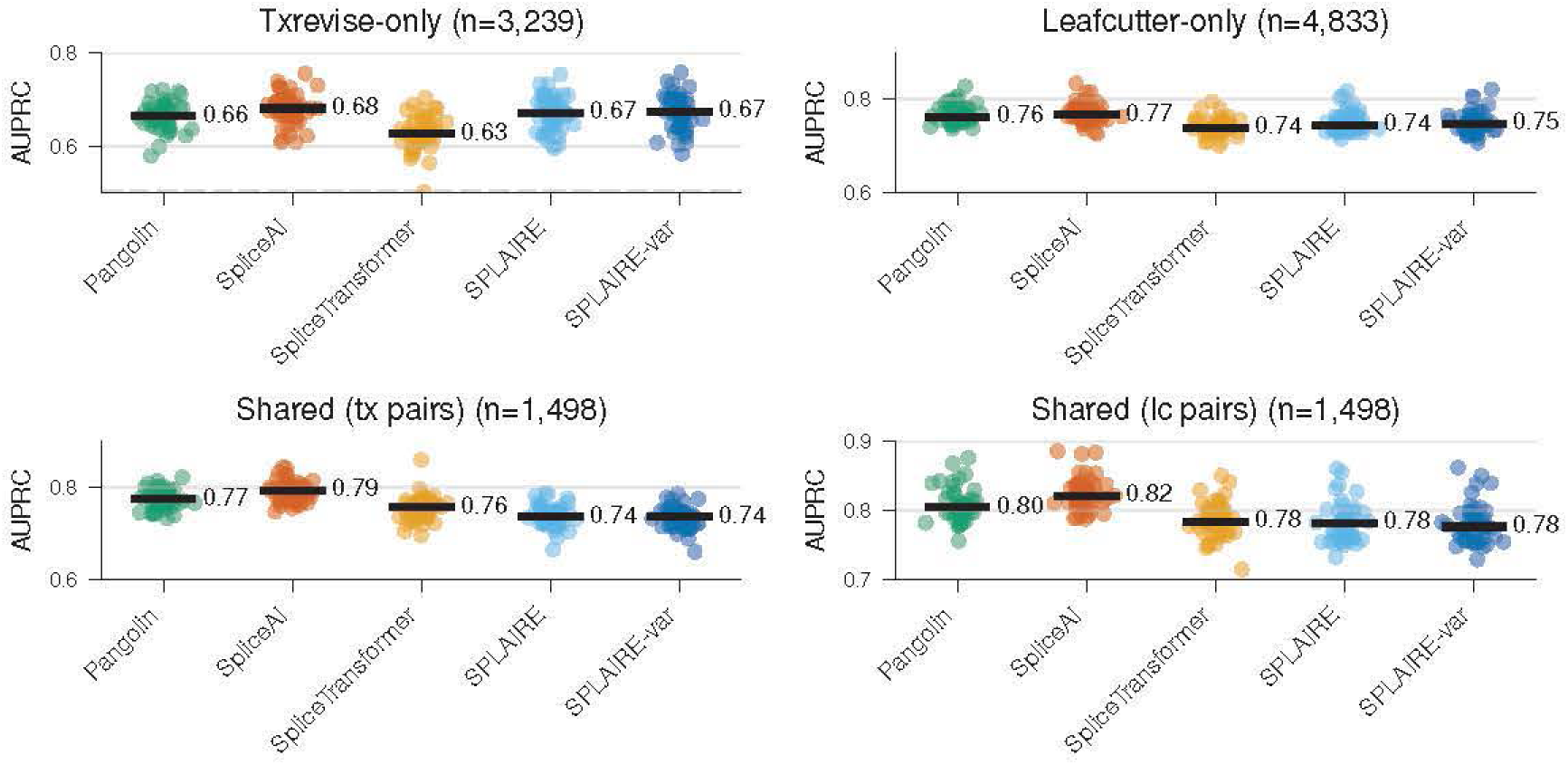
sQTL classification by dataset overlap. (**a**) Per-tissue AUPRC for variants detected as sQTLs only by txrevise (*n* = 3,164 unique variants), evaluated with txrevise-matched negatives. (**b**) Per-tissue AUPRC for variants detected only by leafcutter (*n* = 4,857 unique variants), evaluated with leafcutter-matched negatives. (**c,d**) Per-tissue AUPRC for the same *n* = 1,474 variants detected by both methods, evaluated with txrevise-matched negatives (c) and leafcutter-matched negatives (d).

**Figure S10.**
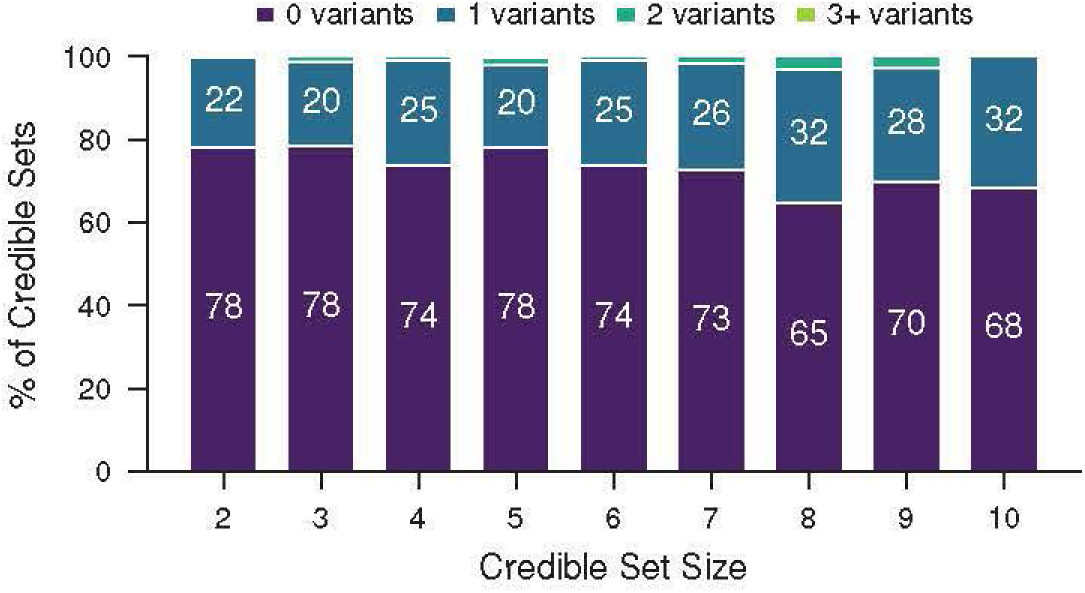
Unresolved credible-set variant counts at score threshold 0.2. Same analysis as Figure 6d but using a threshold of 0.2 in place of the per-tissue Youden’s J threshold. Bars show the percentage of unresolved credible sets (maximum PIP *<* 0.9, 2–10 variants per set) in which SPLAIRE calls 0, 1, 2, or 3+ variants positive, stratified by credible-set size (*n* = 7,402 credible sets across 49 GTEx tissues).

**Figure S11.**
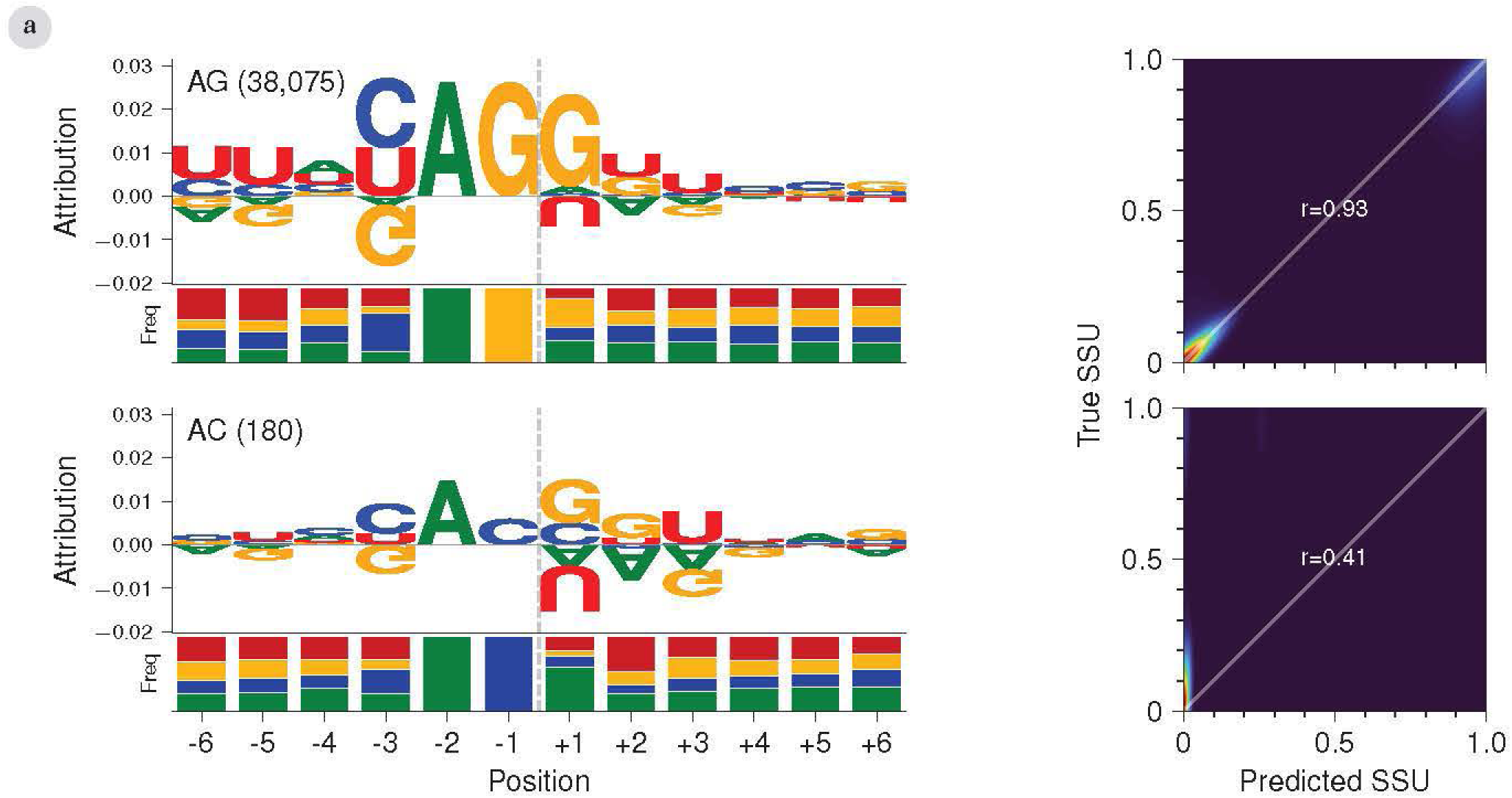
Per-position attribution profiles stratified by acceptor dinucleotide. Acceptor splice sites grouped by the dinucleotide at the last two intronic positions (2/ 1 relative to the intron-exon boundary): AG and AC. For each group, the left column shows the attribution logo from the regression output (top) with a stacked nucleotide frequency bar (bottom) at each position (frequency bar colors match logo letter colors: A green, C blue, G yellow, U red); the right column shows the predicted versus true SSU density with Pearson *r* for that dinucleotide group. Mirrors Figure 7b for acceptor rather than donor splice sites.

**Figure S12.**
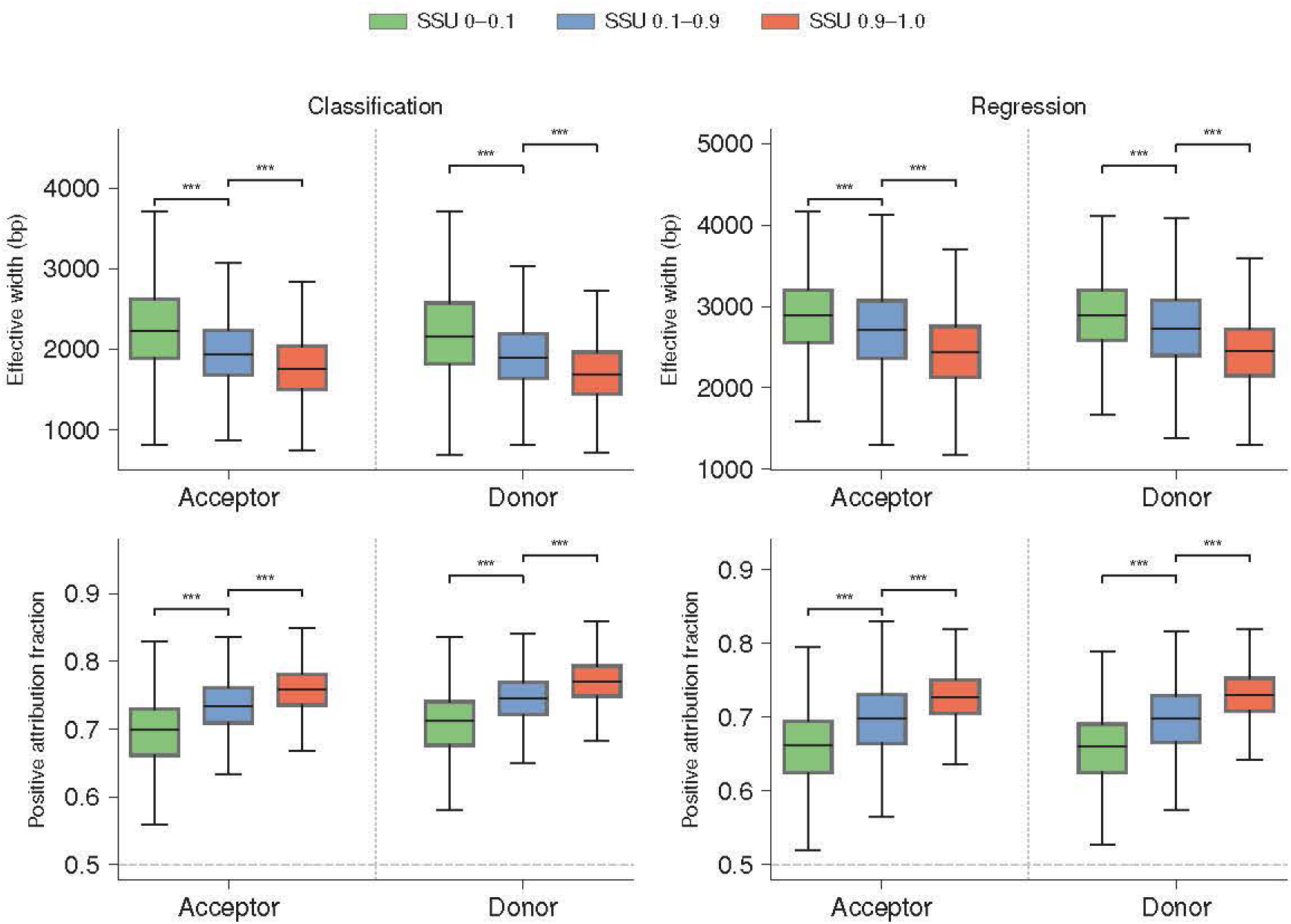
Attribution spread metrics stratified by splice type and SSU. Two metrics quantify the spatial distribution of DeepLIFT-SHAP attributions across the 10,001 bp input sequence, shown for the classification head (left column) and regression head (right column). *Effective width* (top row) is the minimum number of positions capturing 90% of total absolute attribution. *Positive attribution fraction* (bottom row) is the fraction of total attribution magnitude from activating (positive) positions. Dashed line marks 0.5. Splice sites are split by type (acceptor, donor) and true SSU bin, with 0.9–1.0 (red), 0.1–0.9 (blue), and 0–0.1 (green). Significance brackets show pairwise Mann–Whitney *U* tests (Holm–Bonferroni corrected; ∗ *p <* 0.05, ∗∗ *p <* 0.01, ∗ ∗ ∗ *p <* 0.001). *n* = 38,456 acceptor and 34,081 donor splice sites.

**Figure S13.**
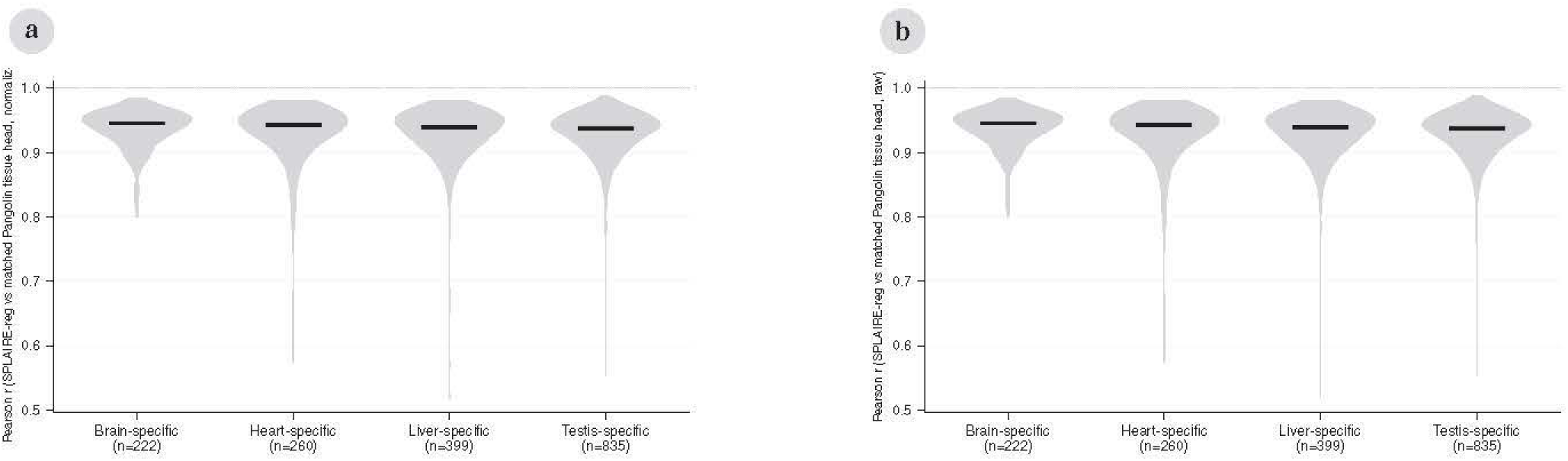
SPLAIRE versus matched Pangolin tissue-head attribution agreement at tissue-specific splice sites. Pairwise Pearson *r* per site between SPLAIRE-reg DeepLIFT-SHAP attributions and the matched-tissue Pangolin DeepLIFT-SHAP attributions, computed on the full 10,001 bp input sequence. Each violin pools the per-site Pearson *r* across all sites where the labeled tissue is the outlier (e.g., Brain-specific = SPLAIRE-reg vs Pangolin-brain at brain-outlier sites). (**a**) L1-normalized attributions. (**b**) Raw attributions.

**Figure S14.**
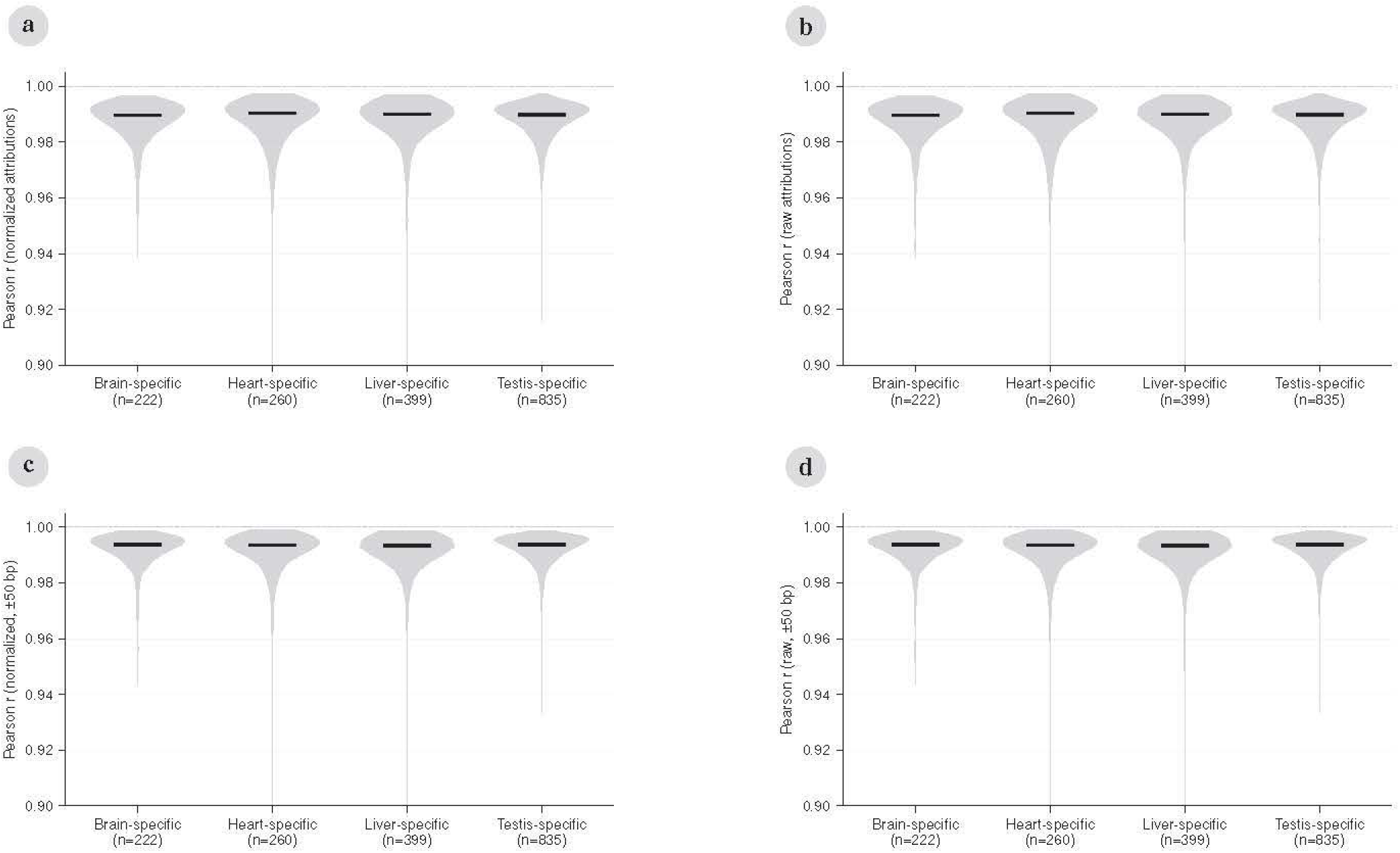
Cross-model attribution agreement at tissue-specific splice sites. Pairwise Pearson *r* between the target Pangolin tissue head and each of the three non-target Pangolin tissue heads, computed from DeepLIFT-SHAP attributions. Each violin pools three model pairs per target tissue (e.g., brain vs heart, brain vs liver, brain vs testis at brain-specific sites). Top row (**a**,**b**) shows correlation computed on the full 10,001 bp input sequence. Bottom row (**c**,**d**) is restricted to the 101 positions surrounding the splice site (±50 bp). (**a**) L1-normalized attributions, full window. (**b**) Raw attributions, full window. (**c**) L1-normalized attributions, ±50 bp window. (**d**) Raw attributions, ±50 bp window.

**Table 1.** MAE on neutral reporter variants near splice sites. Mean absolute error (MAE) on neutral variants (measured ΔPSI within 1 SD of the mean) at proximal splice site (3 exonic and 8 intronic nucleotides adjacent to each splice junction) and intronic positions. Counts in parentheses are the number of neutral variants in each subset.

| Model | Vex-seq | Vex-seq | MFASS | MFASS |
| --- | --- | --- | --- | --- |
|  | Splice site (166) | Intron (742) | Splice site (2,626) | Intron (10,797) |
| Pangolin (cls) | 0.096 | 0.044 | 0.069 | 0.051 |
| SpliceAI (cls) | 0.093 | 0.044 | 0.071 | 0.051 |
| SpliceTransformer (cls) | 0.094 | 0.041 | 0.065 | **0.050** |
| SPLAIRE | 0.083 | 0.039 | **0.064** | 0.050 |
| SPLAIRE-var | **0.082** | **0.038** | 0.064 | 0.050 |

**Table 2.** SSU calibration. OLS slope *m* and intercept *b* of true SSU regressed on predicted SSU for each model and tissue. Perfect calibration corresponds to *m* = 1*, b* = 0. Classification rows use the splice-probability output. Regression rows use the SSU output (SpliceAI has classification only). Bold indicates slope closest to 1 within each output type.

| Model | Output | HAEC | Lung | Testis | Brain cortex | Whole blood |
| --- | --- | --- | --- | --- | --- | --- |
| SpliceAI | Cla | **1.00 (-0.05)** | 0.92 (-0.05) | 0.89 (-0.02) | 0.96 (-0.05) | 0.93 (-0.04) |
| SPLAIRE | Cla | 1.08 (-0.24) | **0.99 (-0.21)** | 0.89 (-0.13) | 1.05 (-0.22) | 1.03 (-0.21) |
| Pangolin | Cla | 1.07 (-0.02) | 0.99 (-0.01) | **1.00 (-0.01)** | **1.03 (-0.02)** | **0.99 (-0.01)** |
| SpliceTransformer | Cla | 0.95 (-0.06) | 0.88 (-0.05) | 0.84 (-0.04) | 0.93 (-0.06) | 0.91 (-0.05) |
| SPLAIRE | Reg | **0.96 (+0.04)** | 0.87 (+0.04) | 0.88 (+0.04) | 0.90 (+0.05) | 0.88 (+0.05) |
| Pangolin | Reg | 1.10 (+0.07) | **1.01 (+0.08)** | 1.06 (+0.06) | **1.04 (+0.09)** | **1.00 (+0.09)** |
| SpliceTransformer | Reg | 1.11 (-0.03) | 1.02 (-0.02) | **1.03 (-0.02)** | 1.07 (-0.02) | 1.03 (-0.02) |

**Table 3.**
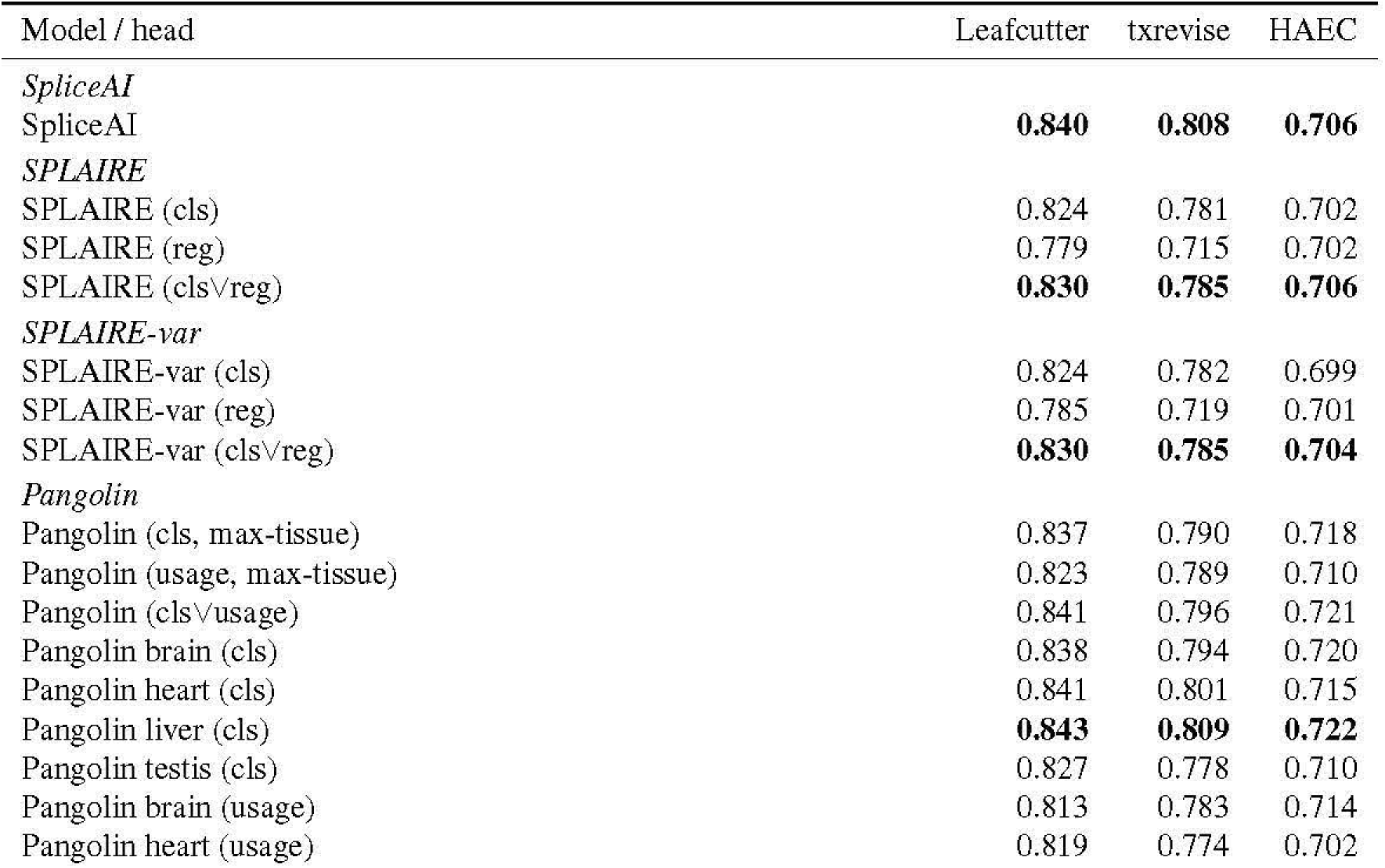
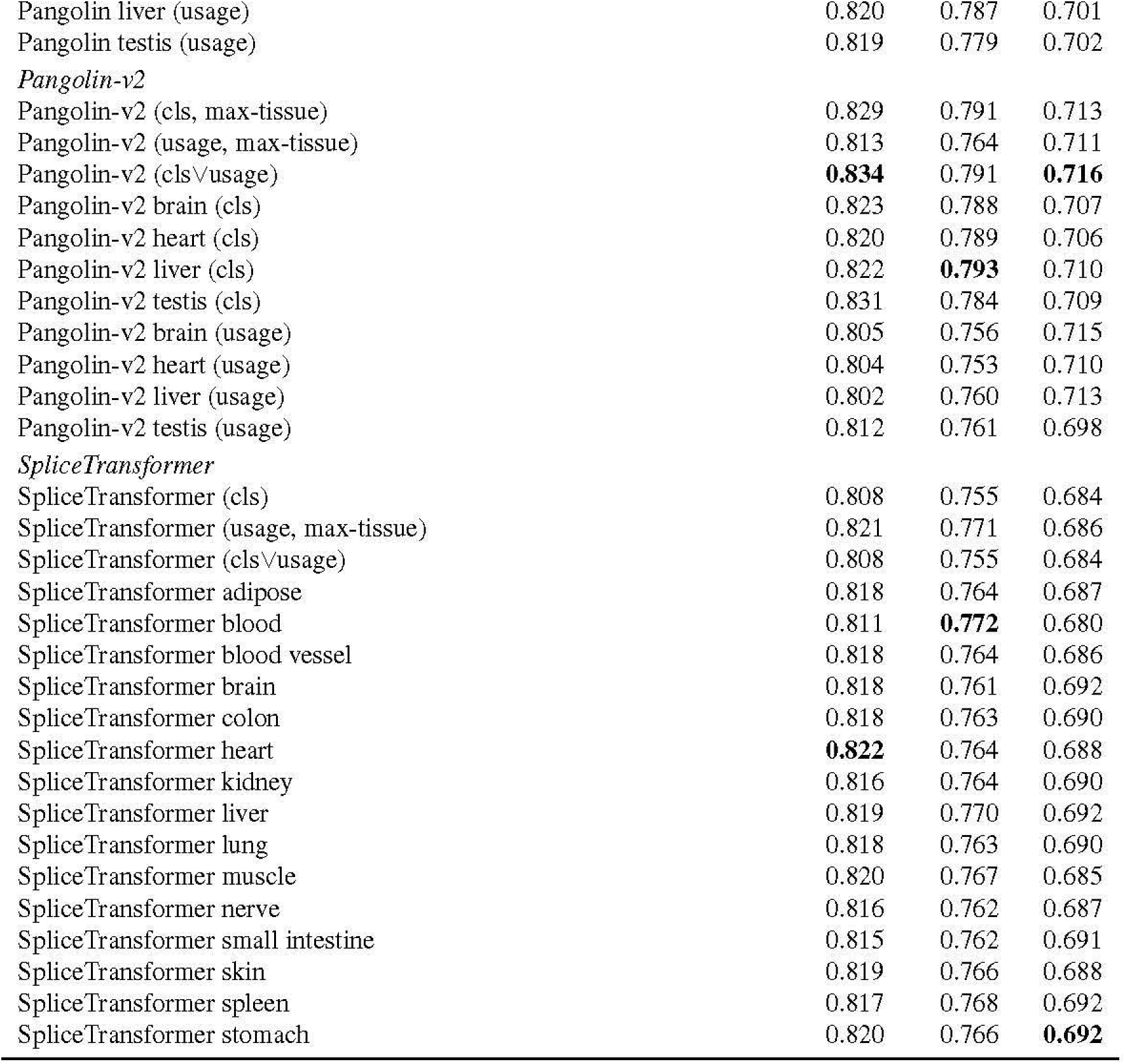
sQTL variant classification extended results. For each model and delta-score formulation, the per-variant score is the maximum absolute delta in the ±2,000 bp window. AUPRC is computed per tissue for the multi-tissue datasets and the median across tissues is reported. HAEC is a single-tissue benchmark.

**Table 4.** Regression metrics by annotation status. Values are means across held-out individuals. *n* is the median number of splice sites per held-out individual. Bold indicates best model per metric within each tissue and annotation subgroup.

| Tissue | Annot. | Model | $n$ | Pearson $r$ | Spearman $\rho$ | $R^2$ | MSE | MAE |
| --- | --- | --- | --- | --- | --- | --- | --- | --- |
| HAEC | Annotated | SPLAIRE | 29,344 | **0.889** | **0.629** | **0.770** | **0.031** | **0.019** |
|  |  | SpliceAI |  | 0.831 | 0.595 | 0.688 | 0.042 | 0.023 |
|  |  | Pangolin |  | 0.852 | 0.572 | 0.539 | 0.062 | 0.122 |
|  |  | SpliceTransformer |  | 0.802 | 0.507 | 0.600 | 0.054 | 0.107 |
|  | Novel | SPLAIRE | 16,918 | **0.110** | **0.147** | -1.543 | 0.009 | **0.002** |
|  |  | SpliceAI |  | 0.095 | 0.107 | -13.705 | 0.051 | 0.043 |
|  |  | Pangolin |  | 0.088 | 0.107 | **-0.537** | **0.005** | 0.002 |
|  |  | SpliceTransformer |  | 0.077 | 0.093 | -10.525 | 0.040 | 0.032 |
| Lung | Annotated | SPLAIRE | 29,513 | **0.823** | **0.608** | **0.626** | **0.049** | **0.040** |
|  |  | SpliceAI |  | 0.770 | 0.585 | 0.531 | 0.062 | 0.061 |
|  |  | Pangolin |  | 0.791 | 0.566 | 0.523 | 0.063 | 0.116 |
|  |  | SpliceTransformer |  | 0.737 | 0.505 | 0.535 | 0.061 | 0.113 |
|  | Novel | SPLAIRE | 14,620 | 0.101 | **0.142** | -2.576 | 0.009 | **0.002** |
|  |  | SpliceAI |  | **0.138** | 0.136 | -20.288 | 0.056 | 0.052 |
|  |  | Pangolin |  | 0.115 | 0.136 | **-0.815** | **0.005** | 0.003 |
|  |  | SpliceTransformer |  | 0.132 | 0.125 | -14.184 | 0.040 | 0.037 |
| Testis | Annotated | SPLAIRE | 36,062 | **0.826** | **0.651** | **0.626** | **0.050** | **0.039** |
|  |  | SpliceAI |  | 0.772 | 0.608 | 0.550 | 0.060 | 0.060 |
|  |  | Pangolin |  | 0.789 | 0.622 | 0.444 | 0.074 | 0.122 |
|  |  | SpliceTransformer |  | 0.741 | 0.566 | 0.519 | 0.064 | 0.115 |
|  | Novel | SPLAIRE | 36,889 | 0.099 | 0.191 | -1.042 | 0.009 | **0.002** |
|  |  | SpliceAI |  | 0.134 | 0.172 | -7.183 | 0.037 | 0.030 |
|  |  | Pangolin |  | **0.152** | **0.210** | **-0.248** | **0.006** | 0.003 |
|  |  | SpliceTransformer |  | 0.150 | 0.177 | -5.818 | 0.031 | 0.032 |
| Brain | Annotated | SPLAIRE | 27,874 | **0.841** | **0.627** | **0.678** | **0.042** | **0.027** |
|  |  | SpliceAI |  | 0.784 | 0.581 | 0.587 | 0.054 | 0.041 |
|  |  | Pangolin |  | 0.815 | 0.582 | 0.518 | 0.063 | 0.121 |
|  |  | SpliceTransformer |  | 0.760 | 0.514 | 0.558 | 0.058 | 0.113 |
|  | Novel | SPLAIRE | 10,779 | 0.084 | 0.138 | -2.271 | 0.012 | **0.002** |
|  |  | SpliceAI |  | 0.105 | 0.119 | -16.974 | 0.067 | 0.062 |
|  |  | Pangolin |  | 0.097 | **0.140** | **-0.801** | **0.007** | 0.003 |
|  |  | SpliceTransformer |  | **0.143** | 0.131 | -10.980 | 0.045 | 0.049 |
| Blood | Annotated | SPLAIRE | 21,986 | **0.831** | **0.594** | **0.656** | **0.045** | **0.031** |
|  |  | SpliceAI |  | 0.773 | 0.561 | 0.557 | 0.057 | 0.048 |
|  |  | Pangolin |  | 0.798 | 0.553 | 0.527 | 0.061 | 0.114 |
|  |  | SpliceTransformer |  | 0.744 | 0.490 | 0.545 | 0.059 | 0.107 |
|  | Novel | SPLAIRE | 9,093 | 0.072 | **0.135** | -2.399 | 0.011 | **0.002** |
|  |  | SpliceAI |  | 0.119 | 0.125 | -16.642 | 0.057 | 0.052 |
|  |  | Pangolin |  | 0.106 | 0.127 | **-0.677** | **0.006** | 0.003 |
|  |  | SpliceTransformer |  | **0.139** | 0.115 | -11.622 | 0.041 | 0.040 |

**Table 5.**
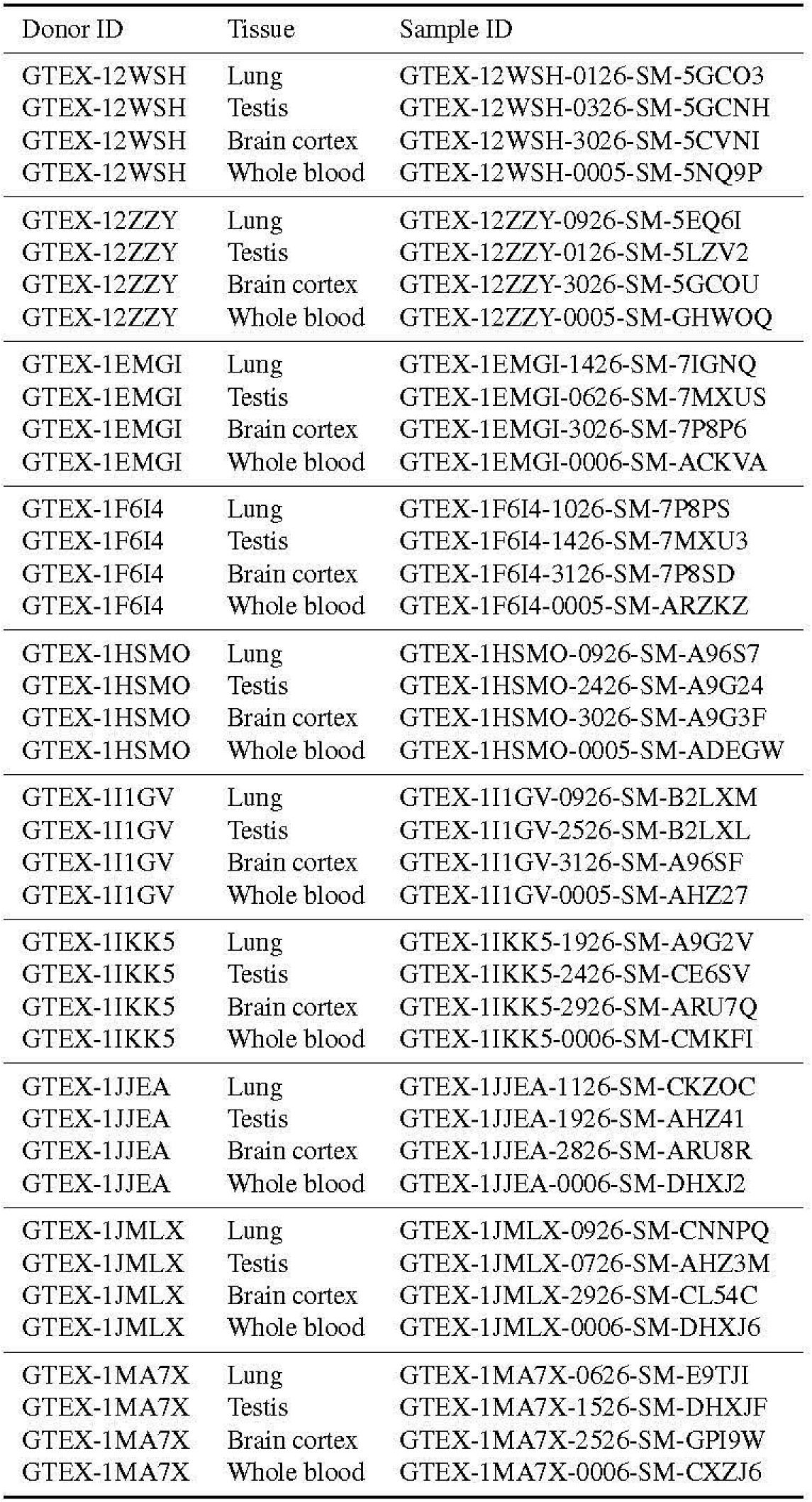
GTEx v10 donor and sample IDs used for held-out evaluation. Ten donors with paired RNA-seq and Whole Genome Sequencing data in lung, testis, brain cortex, and whole blood were selected from the GTEx v10 release.

**Table 6.** Classification metrics (splice site identification, all sites). Values are means across held-out individuals. Top block: classification outputs. Bottom block: regression / usage outputs scored as splice-site classifiers (the regression head’s score is used directly as the classifier score). Bold indicates best per column.

| Head | AUPRC |  |  |  |  |  |  | Top-k |  |  |  |  |  |  |
| --- | --- | --- | --- | --- | --- | --- | --- | --- | --- | --- | --- | --- | --- | --- |
|  | HAEC | Lung | Testis | Brain cortex | Whole blood | GENCODE | MANE | HAEC | Lung | Testis | Brain cortex | Whole blood | GENCODE | MANE |
| SPLAIRE (cls) | **0.753** | **0.762** | 0.679 | 0.771 | 0.735 | 0.947 | **0.948** | 0.685 | 0.694 | 0.618 | 0.713 | 0.698 | 0.888 | **0.871** |
| SPLAIRE-var (cls) | 0.753 | 0.761 | 0.677 | **0.771** | **0.736** | 0.945 | 0.946 | **0.685** | 0.695 | 0.616 | 0.715 | 0.699 | 0.883 | 0.871 |
| SpliceAI (cls) | 0.723 | 0.757 | **0.684** | 0.762 | 0.728 | 0.962 | 0.868 | 0.678 | **0.703** | **0.627** | **0.723** | **0.708** | 0.918 | 0.856 |
| Pangolin (cls, avg) | 0.683 | 0.705 | 0.630 | 0.723 | 0.676 | 0.947 | 0.863 | 0.624 | 0.655 | 0.570 | 0.683 | 0.666 | 0.897 | 0.850 |
| Pangolin (cls, heart) | 0.688 | 0.708 | 0.620 | 0.727 | 0.679 | 0.938 | 0.858 | 0.628 | 0.658 | 0.566 | 0.683 | 0.667 | 0.884 | 0.840 |
| Pangolin (cls, liver) | 0.689 | 0.710 | 0.624 | 0.724 | 0.681 | 0.939 | 0.856 | 0.632 | 0.661 | 0.568 | 0.682 | 0.669 | 0.884 | 0.840 |
| Pangolin (cls, brain) | 0.683 | 0.708 | 0.621 | 0.728 | 0.676 | 0.943 | 0.861 | 0.626 | 0.656 | 0.568 | 0.684 | 0.665 | 0.891 | 0.843 |
| Pangolin (cls, testis) | 0.658 | 0.677 | 0.619 | 0.691 | 0.639 | 0.944 | 0.861 | 0.604 | 0.636 | 0.558 | 0.663 | 0.652 | 0.891 | 0.844 |
| SpliceTransformer (cls) | 0.658 | 0.685 | 0.606 | 0.709 | 0.680 | 0.942 | 0.852 | 0.604 | 0.634 | 0.545 | 0.664 | 0.660 | 0.891 | 0.835 |
| SPLAIRE (reg) | 0.643 | 0.674 | 0.586 | 0.678 | 0.625 | **0.968** | 0.879 | 0.613 | 0.645 | 0.553 | 0.669 | 0.658 | **0.936** | 0.866 |
| SPLAIRE-var (reg) | 0.648 | 0.676 | 0.586 | 0.682 | 0.627 | 0.968 | 0.879 | 0.617 | 0.647 | 0.549 | 0.671 | 0.658 | 0.936 | 0.865 |
| Pangolin (reg, avg) | 0.652 | 0.681 | 0.598 | 0.700 | 0.657 | 0.962 | 0.879 | 0.603 | 0.637 | 0.545 | 0.666 | 0.656 | 0.918 | 0.865 |
| Pangolin (reg, heart) | 0.658 | 0.685 | 0.592 | 0.704 | 0.662 | 0.952 | 0.873 | 0.608 | 0.640 | 0.543 | 0.667 | 0.656 | 0.900 | 0.851 |
| Pangolin (reg, liver) | 0.659 | 0.686 | 0.594 | 0.703 | 0.664 | 0.953 | 0.873 | 0.608 | 0.641 | 0.545 | 0.667 | 0.657 | 0.905 | 0.855 |
| Pangolin (reg, brain) | 0.652 | 0.682 | 0.591 | 0.701 | 0.655 | 0.957 | 0.878 | 0.604 | 0.638 | 0.543 | 0.669 | 0.657 | 0.909 | 0.854 |
| Pangolin (reg, testis) | 0.631 | 0.658 | 0.588 | 0.673 | 0.623 | 0.962 | 0.878 | 0.591 | 0.627 | 0.535 | 0.657 | 0.649 | 0.920 | 0.863 |
| SpliceTransformer (reg, avg) | 0.635 | 0.649 | 0.558 | 0.674 | 0.631 | 0.891 | 0.812 | 0.594 | 0.620 | 0.516 | 0.651 | 0.639 | 0.820 | 0.780 |
| SpliceTransformer (reg, adipose) | 0.634 | 0.647 | 0.556 | 0.672 | 0.627 | 0.887 | 0.809 | 0.594 | 0.618 | 0.515 | 0.649 | 0.638 | 0.815 | 0.775 |
| SpliceTransformer (reg, blood) | 0.636 | 0.649 | 0.559 | 0.673 | 0.636 | 0.875 | 0.795 | 0.597 | 0.621 | 0.522 | 0.647 | 0.638 | 0.790 | 0.751 |
| SpliceTransformer (reg, blood vessel) | 0.634 | 0.644 | 0.554 | 0.669 | 0.624 | 0.880 | 0.803 | 0.593 | 0.615 | 0.514 | 0.647 | 0.636 | 0.808 | 0.769 |
| SpliceTransformer (reg, brain) | 0.630 | 0.647 | 0.553 | 0.674 | 0.629 | 0.899 | 0.821 | 0.590 | 0.616 | 0.512 | 0.650 | 0.640 | 0.838 | 0.794 |
| SpliceTransformer (reg, colon) | 0.635 | 0.649 | 0.557 | 0.674 | 0.630 | 0.892 | 0.813 | 0.594 | 0.619 | 0.516 | 0.651 | 0.639 | 0.823 | 0.783 |
| SpliceTransformer (reg, heart) | 0.632 | 0.645 | 0.553 | 0.670 | 0.625 | 0.882 | 0.805 | 0.592 | 0.615 | 0.513 | 0.647 | 0.635 | 0.810 | 0.772 |
| SpliceTransformer (reg, kidney) | 0.634 | 0.651 | 0.559 | 0.675 | 0.633 | 0.903 | 0.821 | 0.593 | 0.622 | 0.516 | 0.652 | 0.642 | 0.835 | 0.794 |
| SpliceTransformer (reg, liver) | 0.637 | 0.654 | 0.562 | 0.677 | 0.637 | 0.894 | 0.813 | 0.596 | 0.625 | 0.521 | 0.652 | 0.642 | 0.820 | 0.779 |
| SpliceTransformer (reg, lung) | 0.634 | 0.650 | 0.558 | 0.673 | 0.630 | 0.900 | 0.819 | 0.593 | 0.621 | 0.516 | 0.651 | 0.640 | 0.830 | 0.790 |
| SpliceTransformer (reg, muscle) | 0.630 | 0.640 | 0.549 | 0.665 | 0.620 | 0.867 | 0.792 | 0.591 | 0.610 | 0.510 | 0.641 | 0.630 | 0.793 | 0.755 |
| SpliceTransformer (reg, nerve) | 0.633 | 0.646 | 0.555 | 0.671 | 0.626 | 0.887 | 0.810 | 0.592 | 0.617 | 0.514 | 0.648 | 0.638 | 0.818 | 0.778 |
| SpliceTransformer (reg, small intestine) | 0.634 | 0.650 | 0.558 | 0.674 | 0.632 | 0.898 | 0.818 | 0.593 | 0.622 | 0.516 | 0.651 | 0.640 | 0.829 | 0.788 |
| SpliceTransformer (reg, skin) | 0.635 | 0.647 | 0.557 | 0.672 | 0.627 | 0.886 | 0.808 | 0.594 | 0.618 | 0.516 | 0.648 | 0.638 | 0.815 | 0.774 |
| SpliceTransformer (reg, spleen) | 0.635 | 0.651 | 0.559 | 0.674 | 0.634 | 0.889 | 0.809 | 0.596 | 0.623 | 0.519 | 0.650 | 0.640 | 0.813 | 0.773 |
| SpliceTransformer (reg, stomach) | 0.635 | 0.651 | 0.559 | 0.675 | 0.632 | 0.895 | 0.816 | 0.594 | 0.621 | 0.517 | 0.651 | 0.639 | 0.825 | 0.786 |

**Table 7.** SSU prediction performance. Values are means across held-out individuals. Top block contains regression and usage outputs. Bottom block contains classification outputs scored against measured SSU (the maximum of the acceptor and donor probabilities is used for SPLAIRE, SpliceAI, and SpliceTransformer). Bold indicates best per column.

| Head | Pearson $r$ | | | | | Spearman $\rho$ | | | | | $R^2$ | | | | |
| --- | --- | --- | --- | --- | --- | --- | --- | --- | --- | --- | --- | --- | --- | --- | --- |
|  | HAEC | Lung | Testis | Brain cortex | Whole blood | HAEC | Lung | Testis | Brain cortex | Whole blood | HAEC | Lung | Testis | Brain cortex | Whole blood |
| SPLAIRE (reg) | 0.950 | **0.917** | 0.927 | **0.919** | **0.919** | **0.809** | **0.800** | 0.777 | **0.796** | **0.787** | 0.900 | **0.821** | 0.846 | **0.836** | **0.832** |
| SPLAIRE-var (reg) | **0.950** | 0.916 | 0.926 | 0.919 | 0.919 | 0.808 | 0.798 | 0.775 | 0.792 | 0.784 | **0.900** | 0.819 | 0.844 | 0.835 | 0.831 |
| SpliceAI (cls) | 0.902 | 0.869 | 0.885 | 0.867 | 0.871 | 0.789 | 0.784 | 0.761 | 0.770 | 0.767 | 0.801 | 0.702 | 0.746 | 0.721 | 0.722 |
| Pangolin (reg, avg) | 0.934 | 0.900 | 0.909 | 0.905 | 0.903 | 0.794 | 0.789 | 0.780 | 0.783 | 0.774 | 0.819 | 0.783 | 0.793 | 0.771 | 0.783 |
| Pangolin (reg, heart) | 0.920 | 0.882 | 0.885 | 0.887 | 0.888 | 0.791 | 0.785 | 0.772 | 0.779 | 0.770 | 0.783 | 0.740 | 0.738 | 0.723 | 0.747 |
| Pangolin (reg, liver) | 0.909 | 0.870 | 0.876 | 0.871 | 0.879 | 0.788 | 0.781 | 0.772 | 0.771 | 0.765 | 0.745 | 0.701 | 0.708 | 0.672 | 0.716 |
| Pangolin (reg, brain) | 0.937 | 0.905 | 0.907 | 0.913 | 0.905 | 0.797 | 0.794 | 0.776 | 0.791 | 0.778 | 0.836 | 0.801 | 0.798 | 0.799 | 0.796 |
| Pangolin (reg, testis) | 0.942 | 0.908 | **0.929** | 0.911 | 0.910 | 0.797 | 0.789 | **0.782** | 0.783 | 0.775 | 0.862 | 0.818 | **0.853** | 0.811 | 0.817 |
| SpliceTransformer (reg, avg) | 0.892 | 0.857 | 0.868 | 0.861 | 0.862 | 0.765 | 0.757 | 0.751 | 0.746 | 0.739 | 0.786 | 0.731 | 0.752 | 0.738 | 0.740 |
| SpliceTransformer (reg, adipose) | 0.890 | 0.855 | 0.866 | 0.860 | 0.859 | 0.766 | 0.757 | 0.751 | 0.748 | 0.741 | 0.786 | 0.723 | 0.745 | 0.736 | 0.734 |
| SpliceTransformer (reg, blood) | 0.853 | 0.810 | 0.833 | 0.808 | 0.820 | 0.754 | 0.741 | 0.744 | 0.724 | 0.724 | 0.591 | 0.562 | 0.610 | 0.512 | 0.568 |
| SpliceTransformer (reg, blood vessel) | 0.888 | 0.852 | 0.864 | 0.857 | 0.857 | 0.766 | 0.757 | 0.749 | 0.748 | 0.741 | 0.783 | 0.723 | 0.745 | 0.733 | 0.733 |
| SpliceTransformer (reg, brain) | 0.900 | 0.866 | 0.876 | 0.872 | 0.870 | 0.771 | 0.763 | 0.754 | 0.754 | 0.745 | 0.801 | 0.746 | 0.765 | 0.756 | 0.754 |
| SpliceTransformer (reg, colon) | 0.892 | 0.858 | 0.869 | 0.863 | 0.862 | 0.767 | 0.759 | 0.752 | 0.749 | 0.742 | 0.791 | 0.724 | 0.747 | 0.740 | 0.736 |
| SpliceTransformer (reg, heart) | 0.891 | 0.855 | 0.867 | 0.860 | 0.860 | 0.766 | 0.756 | 0.749 | 0.747 | 0.740 | 0.784 | 0.730 | 0.751 | 0.734 | 0.738 |
| SpliceTransformer (reg, kidney) | 0.895 | 0.861 | 0.872 | 0.867 | 0.866 | 0.768 | 0.760 | 0.755 | 0.749 | 0.743 | 0.795 | 0.732 | 0.754 | 0.746 | 0.744 |
| SpliceTransformer (reg, liver) | 0.889 | 0.850 | 0.865 | 0.853 | 0.857 | 0.763 | 0.753 | 0.751 | 0.738 | 0.735 | 0.760 | 0.715 | 0.739 | 0.705 | 0.723 |
| SpliceTransformer (reg, lung) | 0.893 | 0.860 | 0.870 | 0.865 | 0.864 | 0.768 | 0.760 | 0.754 | 0.750 | 0.743 | 0.791 | 0.723 | 0.746 | 0.741 | 0.736 |
| SpliceTransformer (reg, muscle) | 0.883 | 0.845 | 0.858 | 0.849 | 0.850 | 0.762 | 0.752 | 0.745 | 0.744 | 0.737 | 0.763 | 0.712 | 0.733 | 0.710 | 0.719 |
| SpliceTransformer (reg, nerve) | 0.889 | 0.855 | 0.866 | 0.861 | 0.860 | 0.767 | 0.759 | 0.751 | 0.750 | 0.742 | 0.785 | 0.716 | 0.739 | 0.734 | 0.729 |
| SpliceTransformer (reg, small intestine) | 0.891 | 0.857 | 0.868 | 0.863 | 0.862 | 0.767 | 0.759 | 0.753 | 0.747 | 0.741 | 0.787 | 0.716 | 0.740 | 0.735 | 0.730 |
| SpliceTransformer (reg, skin) | 0.890 | 0.855 | 0.867 | 0.860 | 0.860 | 0.766 | 0.757 | 0.750 | 0.748 | 0.741 | 0.788 | 0.725 | 0.748 | 0.737 | 0.736 |
| SpliceTransformer (reg, spleen) | 0.887 | 0.850 | 0.863 | 0.853 | 0.857 | 0.762 | 0.752 | 0.750 | 0.738 | 0.734 | 0.778 | 0.719 | 0.744 | 0.725 | 0.731 |
| SpliceTransformer (reg, stomach) | 0.895 | 0.860 | 0.871 | 0.865 | 0.865 | 0.767 | 0.759 | 0.752 | 0.748 | 0.741 | 0.796 | 0.732 | 0.755 | 0.745 | 0.743 |
| SPLAIRE (cls) | 0.788 | 0.771 | 0.789 | 0.768 | 0.777 | 0.779 | 0.773 | 0.755 | 0.759 | 0.759 | 0.449 | 0.350 | 0.396 | 0.403 | 0.408 |
| SPLAIRE-var (cls) | 0.791 | 0.774 | 0.792 | 0.770 | 0.779 | 0.775 | 0.768 | 0.754 | 0.752 | 0.752 | 0.458 | 0.361 | 0.410 | 0.413 | 0.416 |
| SpliceAI (cls) | 0.902 | 0.869 | 0.885 | 0.867 | 0.871 | 0.789 | 0.784 | 0.761 | 0.770 | 0.767 | 0.801 | 0.702 | 0.746 | 0.721 | 0.722 |
| Pangolin (cls, avg) | 0.932 | 0.897 | 0.912 | 0.902 | 0.899 | 0.785 | 0.779 | 0.775 | 0.767 | 0.761 | 0.865 | 0.800 | 0.831 | 0.812 | 0.805 |
| Pangolin (cls, heart) | 0.928 | 0.890 | 0.900 | 0.895 | 0.893 | 0.783 | 0.775 | 0.766 | 0.763 | 0.757 | 0.853 | 0.792 | 0.808 | 0.799 | 0.797 |
| Pangolin (cls, liver) | 0.925 | 0.885 | 0.899 | 0.889 | 0.890 | 0.780 | 0.771 | 0.766 | 0.756 | 0.753 | 0.845 | 0.783 | 0.805 | 0.786 | 0.791 |
| Pangolin (cls, brain) | 0.931 | 0.897 | 0.908 | 0.902 | 0.897 | 0.785 | 0.780 | 0.769 | 0.770 | 0.761 | 0.864 | 0.795 | 0.823 | 0.811 | 0.800 |
| Pangolin (cls, testis) | 0.919 | 0.883 | 0.902 | 0.886 | 0.887 | 0.787 | 0.780 | 0.778 | 0.769 | 0.765 | 0.834 | 0.745 | 0.785 | 0.766 | 0.762 |
| SpliceTransformer (cls) | 0.875 | 0.842 | 0.846 | 0.847 | 0.850 | 0.772 | 0.771 | 0.755 | 0.762 | 0.754 | 0.730 | 0.609 | 0.617 | 0.651 | 0.651 |

**Table 8.** Classification metrics on the Pangolin test set. AUPRC and top-*k* accuracy for splice site classification across four tissues. Bold indicates best per column.

| AUPRC | Heart | Liver | Brain | Testis |
| --- | --- | --- | --- | --- |
| Pangolin (reg, heart) | 0.860 | 0.808 | 0.864 | 0.750 |
| Pangolin (reg, liver) | 0.861 | 0.814 | 0.863 | 0.755 |
| Pangolin (reg, testis) | 0.854 | 0.810 | 0.863 | 0.783 |
| Pangolin (reg, brain) | 0.858 | 0.807 | 0.867 | 0.752 |
| Pangolin (cls, heart) | 0.865 | 0.813 | 0.872 | 0.759 |
| Pangolin (cls, testis) | 0.859 | 0.815 | 0.871 | **0.799** |
| Pangolin (cls, liver) | 0.865 | 0.818 | 0.870 | 0.763 |
| Pangolin (cls, brain) | 0.866 | 0.813 | 0.877 | 0.763 |
| SPLAIRE (reg) | 0.843 | 0.794 | 0.847 | 0.723 |
| Pangolin (reg, avg) | 0.863 | 0.815 | 0.870 | 0.777 |
| SpliceAI (cls) | 0.838 | 0.784 | 0.841 | 0.720 |
| Pangolin (cls, avg) | **0.869** | **0.821** | **0.878** | 0.790 |
| SPLAIRE-var (reg) | 0.844 | 0.795 | 0.847 | 0.721 |
| SpliceTransformer (reg, adipose) | 0.832 | 0.777 | 0.834 | 0.714 |
| SpliceTransformer (reg, blood) | 0.830 | 0.777 | 0.826 | 0.713 |
| SPLAIRE (cls) | 0.835 | 0.785 | 0.834 | 0.697 |
| SPLAIRE-var (cls) | 0.839 | 0.787 | 0.835 | 0.695 |
| SpliceTransformer (reg, blood vessel) | 0.830 | 0.775 | 0.832 | 0.711 |
| SpliceTransformer (reg, brain) | 0.840 | 0.782 | 0.841 | 0.718 |
| SpliceTransformer (reg, heart) | 0.831 | 0.776 | 0.833 | 0.710 |
| SpliceTransformer (reg, colon) | 0.835 | 0.780 | 0.836 | 0.715 |
| SpliceTransformer (reg, kidney) | 0.840 | 0.784 | 0.840 | 0.723 |
| SpliceTransformer (reg, liver) | 0.838 | 0.783 | 0.836 | 0.721 |
| SpliceTransformer (reg, lung) | 0.837 | 0.782 | 0.838 | 0.720 |
| SpliceTransformer (reg, muscle) | 0.825 | 0.770 | 0.827 | 0.704 |
| SpliceTransformer (reg, small intestine) | 0.837 | 0.782 | 0.837 | 0.718 |
| SpliceTransformer (reg, nerve) | 0.833 | 0.777 | 0.834 | 0.713 |
| SpliceTransformer (reg, spleen) | 0.835 | 0.781 | 0.833 | 0.718 |
| SpliceTransformer (reg, skin) | 0.832 | 0.777 | 0.833 | 0.714 |
| SpliceTransformer (reg, stomach) | 0.836 | 0.780 | 0.837 | 0.717 |
| SpliceTransformer (reg, avg) | 0.836 | 0.780 | 0.836 | 0.716 |
| SpliceTransformer (cls) | 0.832 | 0.774 | 0.834 | 0.718 |

**Table 8.** Top- $k$ (continued)
| Top-$k$ | Heart | Liver | Brain | Testis |
| --- | --- | --- | --- | --- |
| Pangolin (reg, heart) | 0.850 | 0.810 | 0.822 | 0.696 |
| Pangolin (reg, liver) | 0.849 | 0.812 | 0.821 | 0.700 |
| Pangolin (reg, testis) | 0.852 | **0.817** | 0.828 | 0.728 |
| Pangolin (reg, brain) | 0.852 | 0.811 | 0.827 | 0.702 |
| Pangolin (cls, heart) | 0.852 | 0.811 | 0.829 | 0.702 |
| Pangolin (cls, testis) | 0.850 | 0.814 | 0.833 | **0.740** |
| Pangolin (cls, liver) | 0.848 | 0.812 | 0.827 | 0.705 |
| Pangolin (cls, brain) | 0.852 | 0.811 | 0.834 | 0.708 |
| SPLAIRE (reg) | 0.854 | 0.816 | 0.819 | 0.689 |
| Pangolin (reg, avg) | 0.854 | 0.817 | 0.830 | 0.722 |
| SpliceAI (cls) | 0.837 | 0.799 | 0.811 | 0.680 |
| Pangolin (cls, avg) | **0.855** | 0.815 | **0.837** | 0.731 |
| SPLAIRE-var (reg) | 0.854 | 0.816 | 0.818 | 0.684 |
| SpliceTransformer (reg, adipose) | 0.813 | 0.772 | 0.788 | 0.662 |
| SpliceTransformer (reg, blood) | 0.797 | 0.762 | 0.773 | 0.665 |
| SPLAIRE (cls) | 0.823 | 0.784 | 0.792 | 0.645 |
| SPLAIRE-var (cls) | 0.822 | 0.784 | 0.793 | 0.642 |
| SpliceTransformer (reg, blood vessel) | 0.809 | 0.768 | 0.784 | 0.658 |
| SpliceTransformer (reg, brain) | 0.826 | 0.784 | 0.798 | 0.664 |
| SpliceTransformer (reg, heart) | 0.812 | 0.771 | 0.786 | 0.657 |
| SpliceTransformer (reg, colon) | 0.817 | 0.776 | 0.792 | 0.662 |
| SpliceTransformer (reg, kidney) | 0.823 | 0.784 | 0.798 | 0.671 |
| SpliceTransformer (reg, liver) | 0.813 | 0.777 | 0.790 | 0.670 |
| SpliceTransformer (reg, lung) | 0.821 | 0.781 | 0.795 | 0.668 |
| SpliceTransformer (reg, muscle) | 0.801 | 0.761 | 0.777 | 0.651 |
| SpliceTransformer (reg, small intestine) | 0.819 | 0.780 | 0.794 | 0.666 |
| SpliceTransformer (reg, nerve) | 0.814 | 0.773 | 0.789 | 0.660 |
| SpliceTransformer (reg, spleen) | 0.809 | 0.772 | 0.786 | 0.667 |
| SpliceTransformer (reg, skin) | 0.811 | 0.771 | 0.787 | 0.661 |
| SpliceTransformer (reg, stomach) | 0.818 | 0.778 | 0.793 | 0.664 |
| SpliceTransformer (reg, avg) | 0.815 | 0.775 | 0.792 | 0.664 |
| SpliceTransformer (cls) | 0.833 | 0.795 | 0.803 | 0.675 |

**Table 9.**
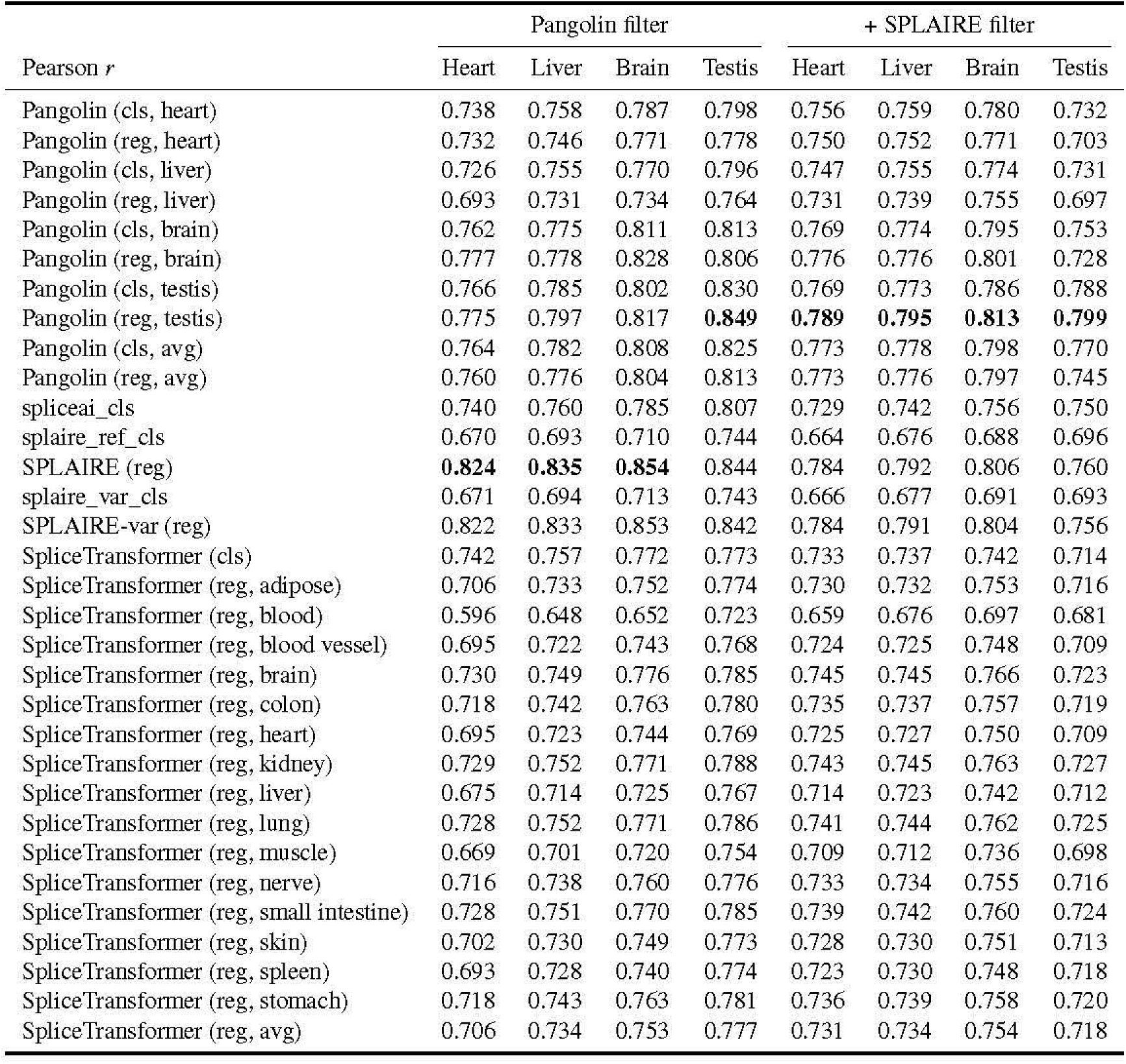
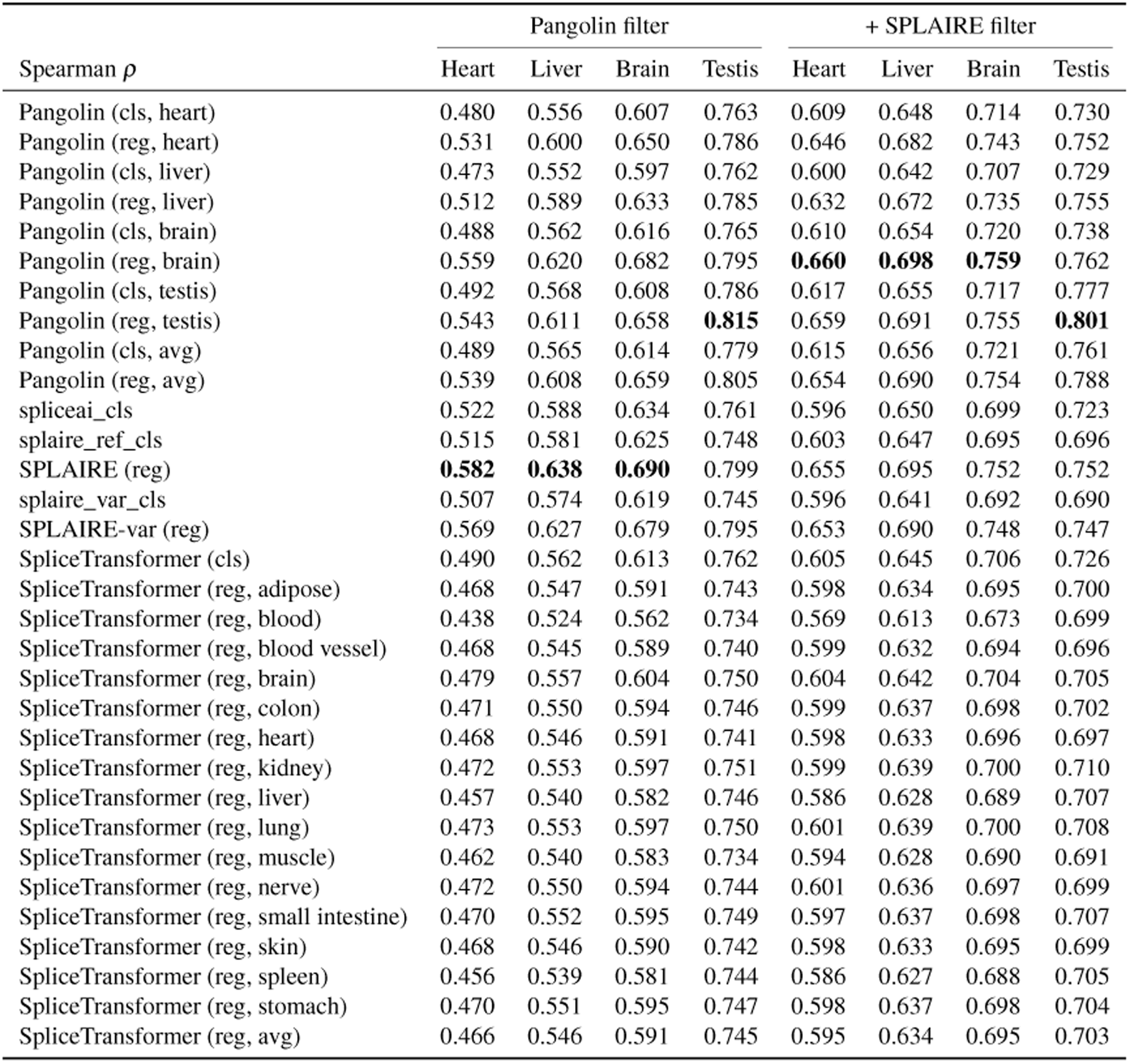
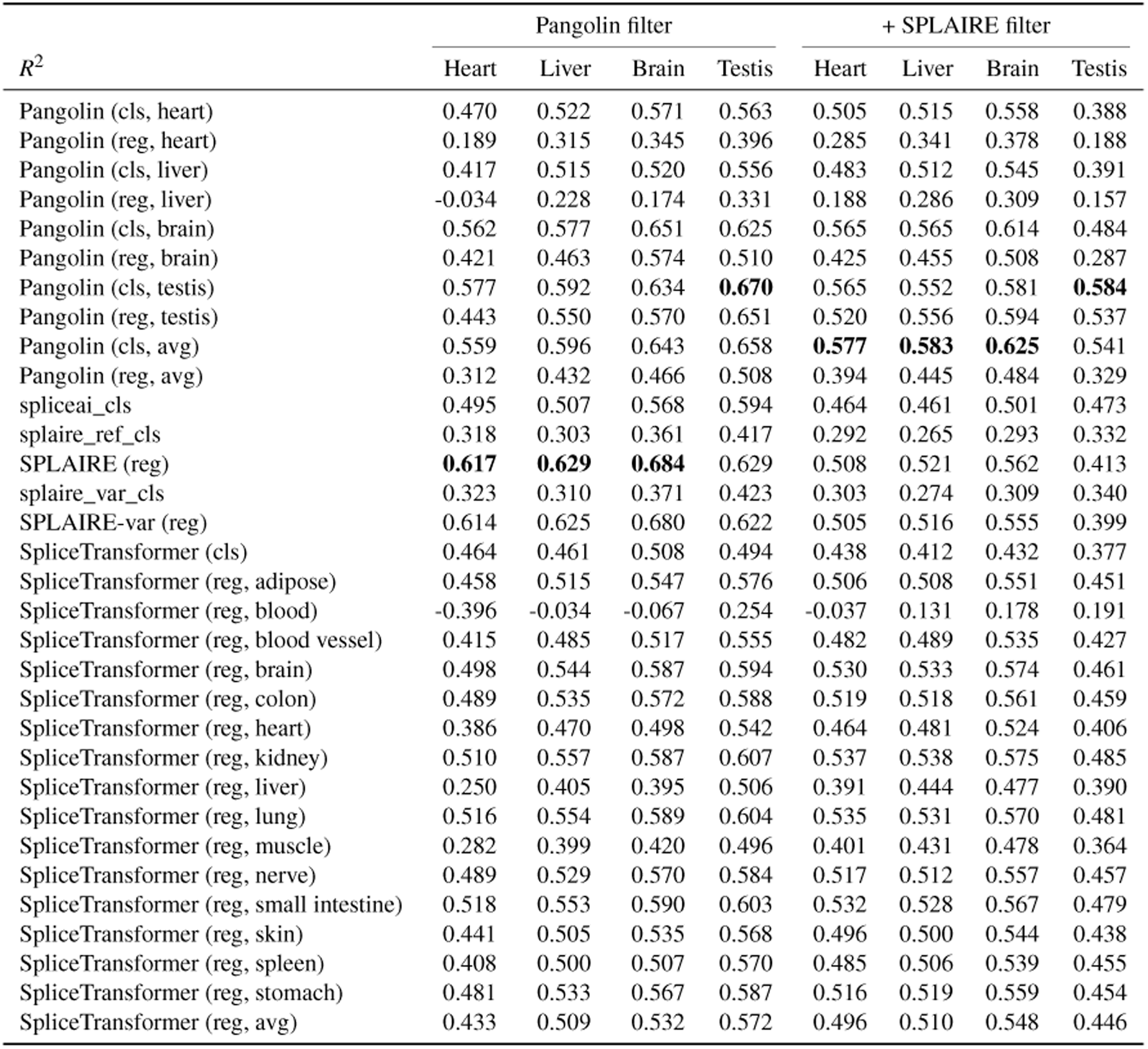
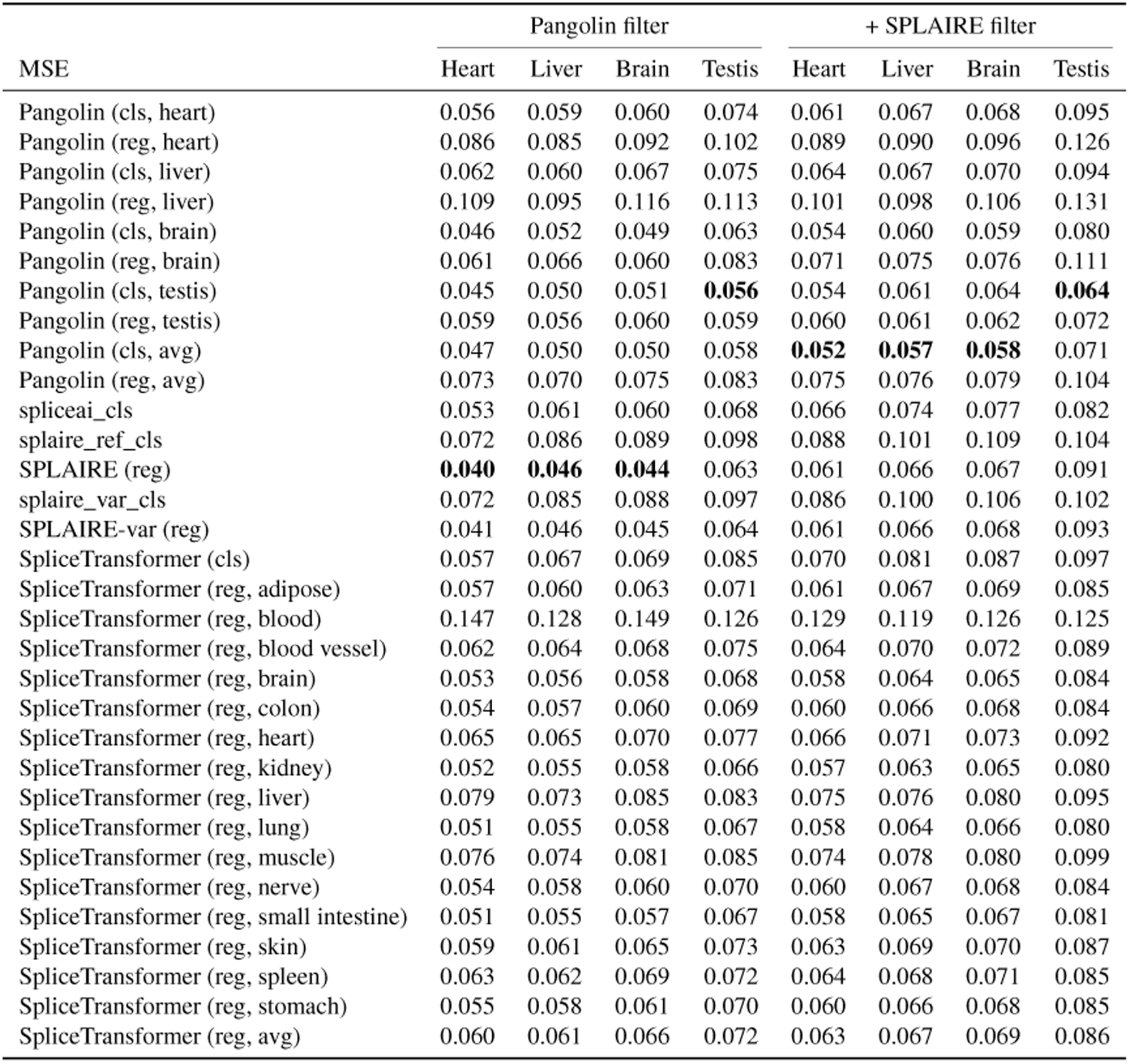
SSU regression metrics on the Pangolin test set. Pearson *r*, Spearman *ρ*, *R*^2^, and MSE for each model across four tissues from the Pangolin test set, reported under Pangolin’s published paralog filter (left) and after additionally applying our SPLAIRE paralog filter (right). Bold indicates best per column within each filter.

**Table 10.** MAE on tissue-specific splice sites on the SPLAIRE test set. Median absolute error on tissue-specific splice sites detected in all five evaluation tissues (HAEC, lung, testis, brain cortex, whole blood) per target tissue (the tissue whose mean SSU deviated most from the cross-tissue mean; *τ* ≥ 0.5 or *τ*_rev_ ≥ 0.7 with cross-tissue SSU range ≥ 0.3; Methods). For Pangolin and SpliceTransformer, the tissue-matched head MAE is shown alongside the lowest MAE among non-matched heads (best alternative). SPLAIRE and SpliceAI do not have tissue-specific output heads.

| Target tissue (n) | SPLAIRE | SpliceAI | Pangolin |  | SpliceTransformer |  |
| --- | --- | --- | --- | --- | --- | --- |
|  | MAE | MAE | matched | best alt | matched | best alt |
| HAEC (3,752) | 0.139 | 0.133 | – | 0.154 | – | 0.154 |
| Lung (3,304) | 0.369 | 0.345 | – | 0.252 | 0.294 | 0.201 |
| Testis (11,688) | 0.412 | 0.394 | 0.323 | 0.288 | – | 0.224 |
| Brain Cortex (2,244) | 0.413 | 0.370 | 0.319 | 0.333 | 0.292 | 0.256 |
| Whole Blood (3,360) | 0.581 | 0.574 | – | 0.391 | 0.302 | 0.402 |

